# Pain-related learning signals in the human insula

**DOI:** 10.1101/2022.01.16.476547

**Authors:** Björn Horing, Christian Büchel

## Abstract

Pain is not only a perceptual phenomenon, but also a preeminent learning signal. In reinforcement learning models, prediction errors (PEs) play a crucial role, i.e. the mismatch between expectation and sensory input. In particular, advanced learning models require the representation of different types of PEs, namely signed PEs (whether more or less pain was expected) to specify the direction of learning, and unsigned PEs (the absolute deviation from an expectation) to adapt the learning rate. The insula has been shown to play an important role in pain intensity coding and in signaling surprise. However, mainly unsigned PEs could be identified in the anterior insula. It remains an open question whether these PEs are specific to pain, and whether signed PEs are also represented in the insula.

To answer these questions, 47 subjects learned associations of two conditioned stimuli (CS) with four unconditioned stimuli (US; painful heat or loud sound, of one low and one high intensity each) while undergoing functional magnetic resonance imaging (fMRI) and skin conductance response (SCR) measurements. CS-US associations reversed multiple times between intensities and between sensory modalities, generating frequent PEs.

SCRs indicated comparable nonspecific characteristics of the two modalities. fMRI analyses focusing on the insular and opercular cortices contralateral to painful stimulation showed that activation in the anterior insula correlated with unsigned intensity PEs. Importantly, this unsigned PE signal was similar for pain and aversive sounds and also modality PEs, indicating an unspecific aversive surprise signal. Conversely, signed pain intensity PE signals were modality-specific and located in the dorsal posterior insula, an area previously implicated in pain intensity processing.

Previous studies have identified abnormal insula function and abnormal learning as potential causes of pain chronification. Our findings link these results and suggest one potential mechanism, namely a misrepresentation of learning relevant prediction errors in the insular cortex.

## Introduction

Apart from its role in signaling tissue damage, pain is increasingly considered to be a preeminent teaching signal [1, 2] in the context of reinforcement learning models [3]. For example, delta rule learning models in classical fear conditioning, such as the Rescorla-Wagner model [4], almost exclusively employ pain as unconditioned stimulus (US). In this and similar models, the value of predictive cues (conditioned stimuli, CS) is updated by the difference between the expected and the experienced outcome, i.e. a prediction error (PE). In this case the PE needs to be *signed* and signals the direction of the difference between expectation and event, i.e. whether the outcome is better or worse than expected. In the case of an aversive event like painful stimulation, this is relevant for shaping future behavior. Reinforcement learning particularly relies on these valences, and different neuronal correlates have been reported for aversive compared to appetitive PEs [5–8]. This has important clinical implications, as pathological learning mechanisms [1, 9] have been reported in chronic pain.

However, PEs can also be computed as *unsigned* [10–12]. An unsigned PE simply indicates the presence of an unexpected event regardless of its valence. Unsigned PEs are therefore conceptually related to constructs like surprise or salience, and may contain information concerning the urgency of behavioral change [13]. Computational models of learning can include either type of PE, or both [4,10,14–16] – for example, the Pearce-Hall model incorporates the unsigned PE as a factor to increase the learning rate after highly incongruent (surprising) events [14, 17], whereas a hybrid-model contains both terms [10,17,18].

Previous studies investigating PEs in the context of aversive learning have observed signal changes in the anterior insula related to unsigned PEs [6,12,19–21]. Unfortunately, in many studies, a signed PE signal is non-orthogonal to stimulus expectation, which poses a problem with a short interval between CS and US, and the low temporal resolution of functional magnetic resonance imaging (fMRI). Consequently, these studies were suboptimal to investigate signed PEs.

Granted that unsigned PEs resemble a surprise signal, they could plausibly involve similar regions for all surprising events, independent of the stimulus sensory modality. Crucially, the representation of unsigned pain PEs in the anterior insula [12, 19] raises the question of whether these are specific to pain, or simply related to aversive events.

To further investigate the existence of signed PEs and the modality-specificity of unsigned PEs, as well as the underlying neuronal mechanisms, we used a Pavlovian transreinforcer reversal learning paradigm [22, 23]. This involves two visual stimuli as CS, and two intensities of painful heat or loud sounds as US (for brevity, these are referred to as “pain” and “sound” forthwith). Across sensory modalities, stimuli were chosen to be roughly comparable in salience as indicated by similar skin conductance responses (SCR) [24]. Reversals occurred between US intensity but within US modality (e.g. CS predicting low pain will next predict high pain), or within US intensity but between US modality (e.g. CS predicting loud sound will next predict high pain). Analyses focused on PEs within and across modalities, using advanced surface based analyses of high resolution fMRI together with skin conductance responses.

We expected that SCR resembles unsigned PEs, as SCR is generally considered to reflect arousal-related activation [25–27] and thus the sign of the PE – representing its valence – should not affect it. Concerning fMRI, we expected to replicate previous results [12, 19] showing the representation of unsigned PEs in the anterior insula. More importantly, we expected that this signal occurs independent of the modality of the US (i.e. both for sound and pain). In agreement with this nonspecific response, we also expected modality PEs to be represented in the anterior insula. However, in this case we expected a weaker signal, as the intensity – and thus salience and other general aspects – are intendedly not different between the expected and the received US.

Employing our novel paradigm, we were also in the position to investigate signed intensity PEs. Focusing on pain, we expected them to be either represented as a distinct part of the anterior insula, or within the mid to posterior insula. The former is suggested by inherent differences in salience between the two intensities, the latter by the notion that a signed PE necessitates some form of intensity encoding, which has been observed in the dorsal posterior insula [24,28– 30].

## Results

In two sessions with 64 trials each, 47 subjects learned associations of two conditioned stimuli (fractal pictures; CS) with individually calibrated unconditioned stimuli (US; two painful heat intensities and two loud sound intensities) (Figure 1a, b). In each trial, either CS appeared, followed by symbols of all four US, from which subjects selected the US they expected (Figure 1c). One of the US was then applied. CS/US associations were deterministic, but importantly, associations frequently reversed and had to be relearned over the course of the experiment (Figure 2). Reversals occurred unannounced after a randomized number of trials. Reversals could occur along the modality dimension or the intensity dimension, but not both simultaneously (e.g., no low heat to high sound reversals). See Materials and Methods and Supporting Figure 1 for further details concerning design and protocol.

**Figure 1.**
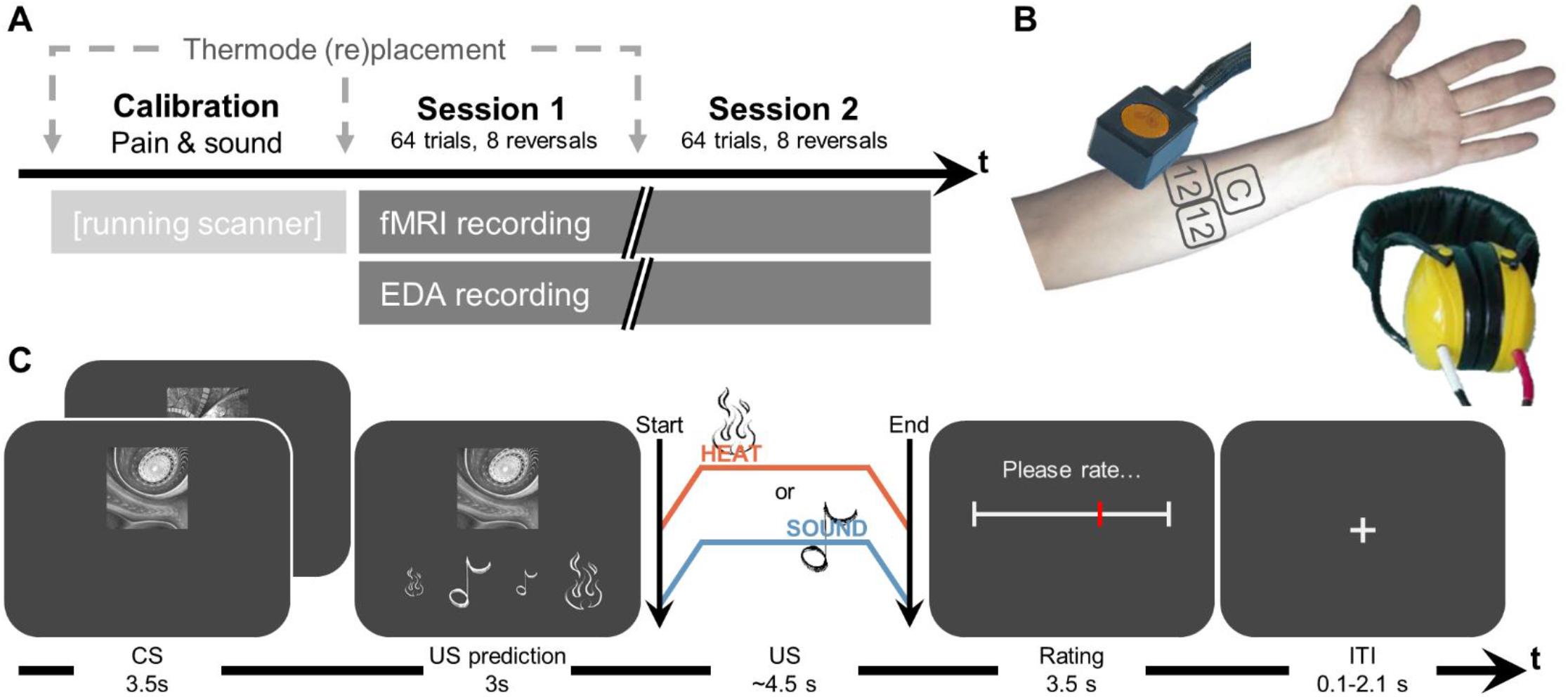
Experimental protocol. **(A)** Overall structure of the experiment. Calibration took ∼15 minutes, each session ∼20 minutes. **(B)** Devices used for heat stimulation (thermode) and sound stimulation (headphones), with standardized locations on the left arm for pain calibration and either of the two experimental sessions. **(C)** Trial structure with associated durations. After displaying CS, subjects were asked to choose which US they expected to follow. The US was then applied and rated in terms of its painfulness (for pain)/unpleasantness (for sound). EDA, electrodermal activity; CS, conditioned stimuli; US, unconditioned stimuli.

**Figure 2.**
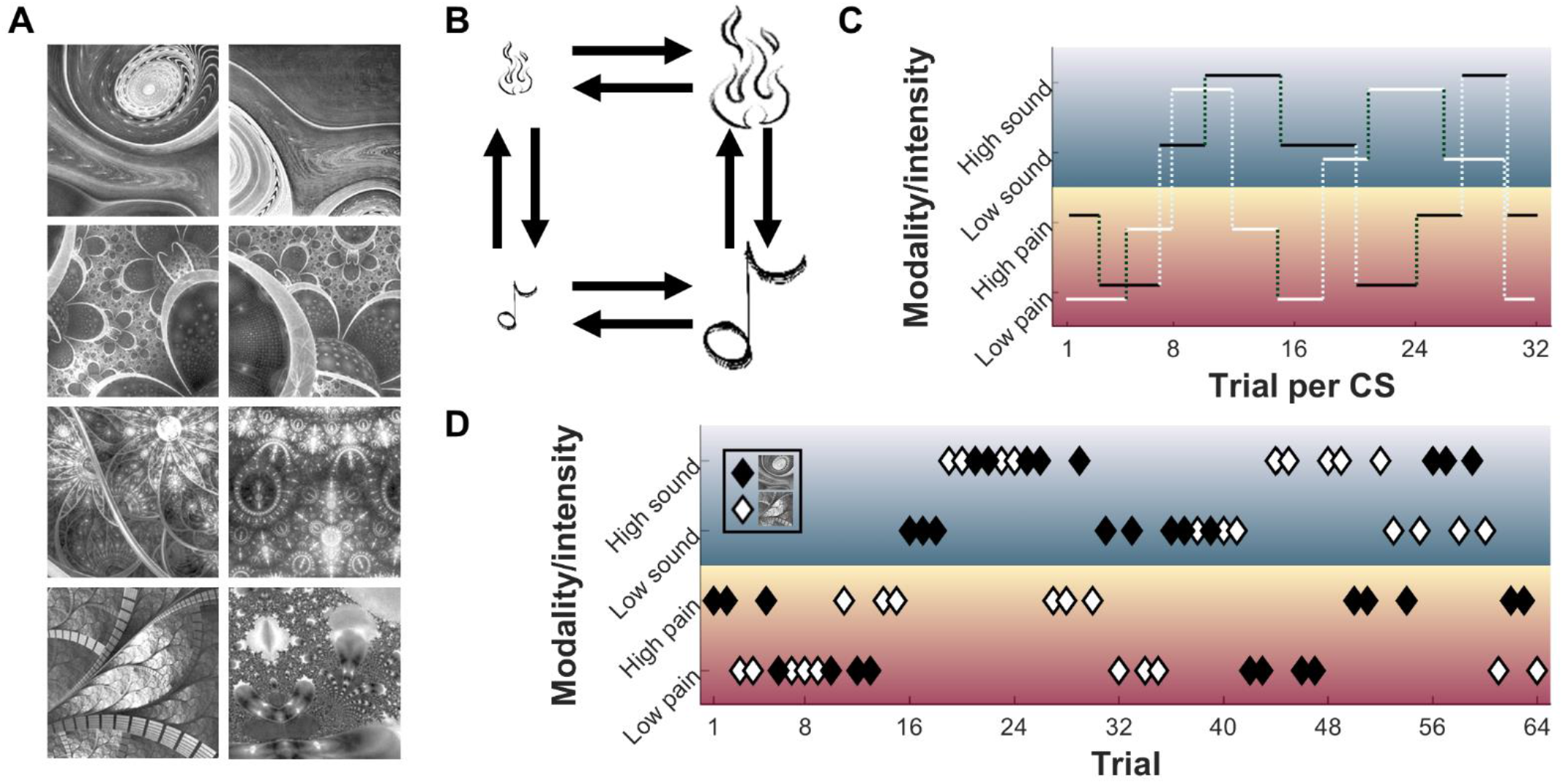
Learning protocol-related aspects of the experiment. **(A)** Set of conditioned stimuli; two were randomly selected for each subject (constraint: stimuli in row 2 could never both be selected due to high similarity). **(B)** Possible US associated with a CS at any particular trial (low pain, high pain, low sound, high sound). Arrows indicate possible reversals; notably, no combined intensity *and* modality (cross)reversals occurred. **(C)** Example for contingencies of CS1 (black solid line) and CS2 (white solid line) for their 32 trials per session each. Vertical dotted lines indicate reversals, with light dotted lines for modality reversals, dark dotted lines for intensity reversals. **<u>(D)</u>** Example for an actual trial sequence of 64 trials with interspersed CS1 (black diamonds) and CS2 (white diamonds), and their associated US (rows). CS, conditioned stimuli; US, unconditioned stimuli.

### Behavioral results: Calibrated stimulus intensities

Calibration yielded temperatures of 44.4±1.2°C for the less painful stimulus (25VAS) and 46.8±1.2°C for the more painful stimulus (75VAS). For sound, calibration yielded 91.7±2.8dBA for the less loud sound (25VAS) and 97.9±3.7dBA for the louder sound (75VAS). Distributions of calibrated stimulus intensities are displayed in Supporting Figure 2a.

### Behavioral results: Stimulus ratings

The first question concerning the behavioral data was whether ratings corresponded to the calibrated intensities (supposed to yield VAS of 25 and 75, respectively). Actual low pain ratings were at 15.4±14.8VAS, high pain ratings at 66.8±21.3VAS; low sound ratings were at 29.2±21.0VAS, high sound ratings at 63.3±19.4VAS (Figure 3a; see Supporting Figure 2b for individual ratings per subject).

**Figure 3.**
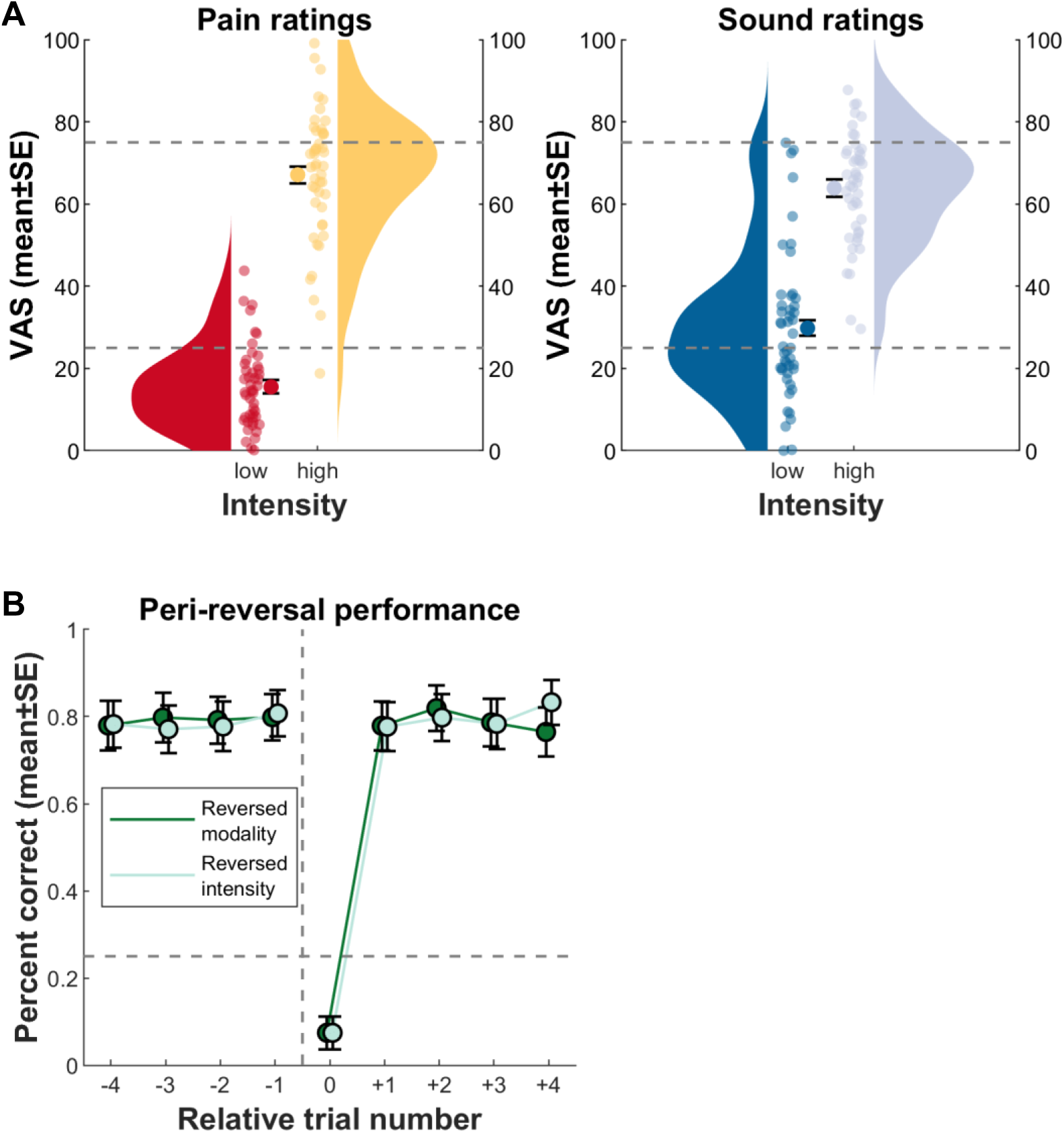
Behavioral results for pain ratings and performance. **(A)** Results for low and high unconditioned pain and sound stimuli; aggregate ratings of all pain and sound trials. Circles with error bars show the mean ± standard errors over all subject means. Subject means are displayed as smaller circles. Violin plots aggregate over subject means. The grey dashed line is the “intended” rating as per calibration (VAS25 for low, VAS75 for high intensities). **(B)** Performance pre and post reversals, aggregated over all subjects. Circles indicate the performance during (peri)reversal trials, first averaged within and then between subjects (mean ± standard errors). The dashed horizontal line marks chance level (25%, i.e. 1 of 4 options). The dashed vertical line indicates contingency reversal, with relative trial number 0 as the reversal trial. Note that no difference arose between trials preceding and following modality versus intensity reversals (also see Figure 2 for aspects concerning contingency reversals). Furthermore, the steep increase in performance after trial number 0 indicates, on average, rapid learning of the new contingency.

### Behavioral results: Learning performance

The next behavioral question was whether the subjects learned the CS/US contingencies. Figure 3b depicts mean performance in predicting the US currently associated with the CS, in relation to the reversals of the association. Combining reversal types and comparing performance at the single trials prior reversal, at reversal, and after reversal, we find pre-reversal performance to be above chance level (t[79] = 13.8, p ≈ 0), at reversal performance below chance (t[79] = -15.9, p ≈ 0), and post-reversal performance back above chance (t[79] = 19.5, p ≈ 0).

### Skin conductance response results

The major question concerning SCR results were whether any differences between the US arose, and how the different PE types would be reflected in this psychophysiological measure of nonspecific characteristics or processes like arousal, salience, or surprise. SCR following sound has a faster onset than that following heat pain stimuli (Figure 4a; see Materials and Methods concerning the different response windows). The average amplitude of pain-related SCR was higher than the average of sound-related SCR, but this difference only showed a trend towards significance (main effect modality, t[4399] = -1.7228, p = 0.08499). Instead, the difference is subsumed by a larger difference between low and high stimuli in the pain modality, as compared to thats in the sound modality (modality*intensity, t[4399] = -2.9739, p = 0.0029567). On average, higher stimuli lead to larger amplitude as well (main effect intensity, t[4399] = 8.2743, p= 1.7 x 10^-16^). Investigating this difference only in correctly predicted trials shows a similar effect on SCR (modality, t[2674] = - 1.4379, p = 0.1506; intensity, t[2674] = 8.0081, p = 2 x 10^-15; modality*intensity, t[2674] = -4.6669, p = 3 x 10^-6) (Supporting Figure 3, Supporting Table 1).

**Figure 4.**
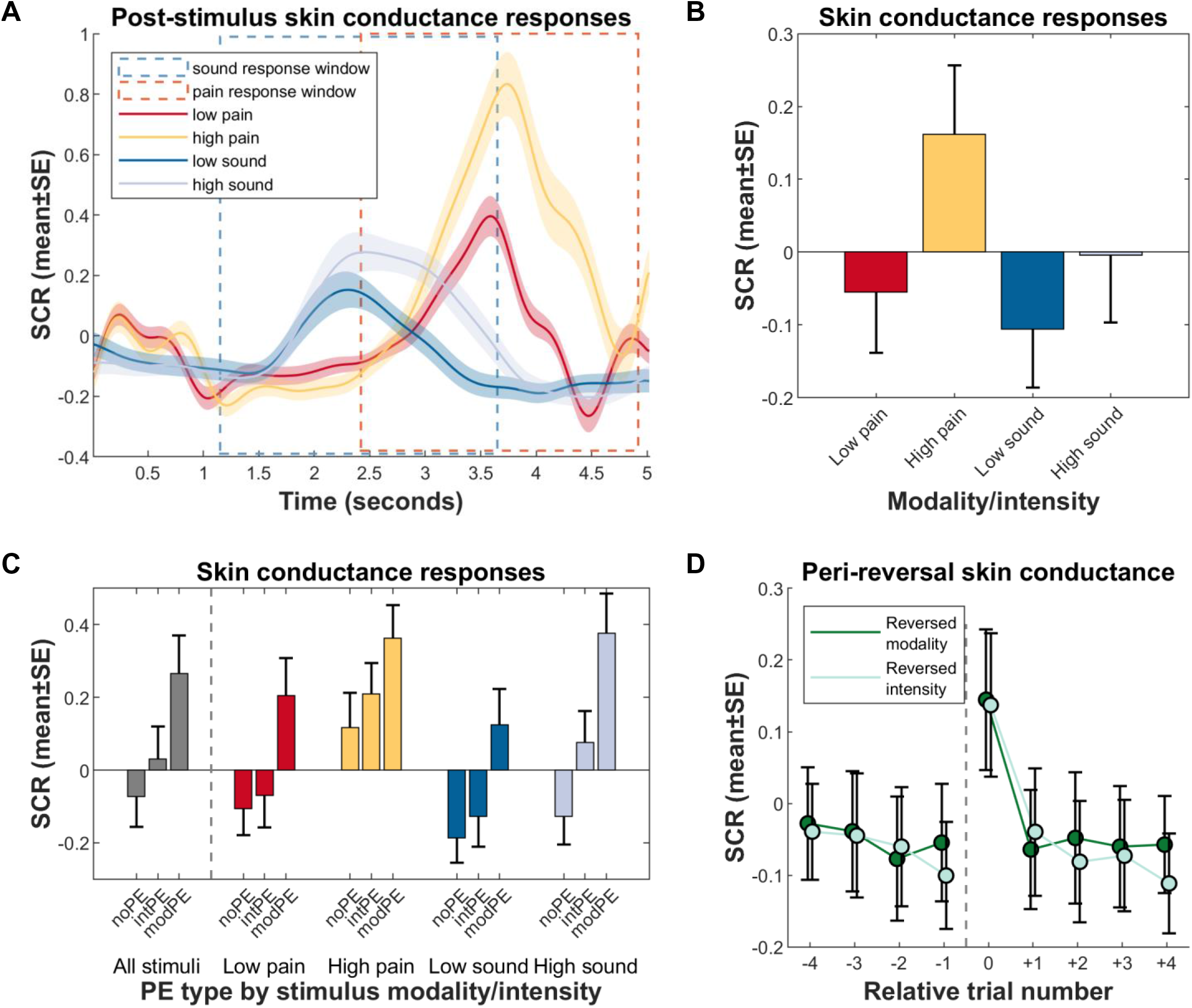
Results from skin conductance response measurements. All plots are based on log- and z-transformed data. **(A)** SCR in relation to unconditioned stimulus onsets, by US modality/intensity. Note the differences in latencies between the two modalities (pain in red/yellow has a later onset, sound in dark blue/light blue earlier), which determined the response windows used for mean SCR calculation in panel b. **(B)** Mean SCR by US, calculated within each modality’s response window. On average, SCR is not significantly different between modalities; differences arise between intensities, and in the interaction of modality and intensity (see text for parameters). **(C)** Mean SCR by US and prediction error type. Over all modalities and intensities, differences arise between each PE type. Within specific modality/intensity combinations, differences between no PEs and intensity PEs only arise in the high sound condition. **(D)** Mean SCR in and around reversal trials. The dashed vertical line indicates contingency reversal, with relative trial number 0 as the reversal trial. SCR rises sharply after reversal, but quickly adapts post reversal to a stable level. SCR, skin conductance response; US, unconditioned stimulus; PE, prediction error; noPE, correct prediction; intPE, intensity prediction error; modPE, modality prediction error.

Further investigating SCR differences following PEs, we first distinguished SCR when subjects correctly predicted the US from trials when either an intensity PE or modality PE was made (Figure 4c). The following statistics include all trials – not just reversals – where an incorrect prediction was made. As shown in the first block (grey bars), over all US and controlling for modality and intensity, SCR following unsigned intensity PEs are larger than those following no PE (intPE>noPE, t[4397] = 4.336, p = 2 x 10^-05^), while SCR following modality PEs are even larger (modPE>noPE, t[4397] = 12.345, p = 2 x 10^-34^; modPE>intPE, t[4397] = 6.398, p = 2 x 10^-10^).

Notably, we performed an adjunct analysis on whether the direction of intensity PEs (i.e. signed intensity PEs) had an impact. We obtained mean SCR differences per subject between no PE and intensity PE trials for each modality and intensity separately, thereby accounting for higher intensity-related base SCRs; next, we contrasted these (now signed) PE-related differences between the low and high intensity. For pain, results indicate no effect (PE-related SCR difference for low pain mean±SE 0.036±0.052, for high pain 0.0922±0.0622, paired t-test t[36] = -0.725, p = 0.4731), while for sound, a more ambiguous yet non-significant result arose (PE-related SCR difference for low sound mean±SE 0.060±0.054, for high sound 0.199±0.054, paired t-test t[35] = -1.931, p = 0.0616).

In four consequent analyses, we investigated differences in SCR following PEs in all US separately, meaning that all intensity PEs are now signed. Results indicate that the intPE>noPE effect of the global analysis is driven by this contrast in the high sound US (light blue bars, t[1119] = 4.732, p = 3 x 10^-6^); it does not reach significance following any other US. Conversely, modality PEs are followed by larger SCR in all US (all modPE>noPE p < 0.001; smallest effect modPE>intPE t[1090] = 2.045, p = 0.041079).

Figure 4d shows the average perireversal trial effect on SCR, over all US. It shows a large increase in SCR during both modality and intensity reversals; note that this analysis does not consider actual subject expectation, just the position related to the reversal trial. SCR is highest during the reversal trial, and rapidly reaches a lower plateau even one trial later. Comparing the pre-reversal trial to immediate post-reversal (trials -1 to +1), SCR is not significantly different if a modality reversal occurred (p = 0.54704); this is also the case if an intensity reversal occurred (p = 0.071164).

### Imaging results

We first obtained an overview of modality-related effects (Figure 5a/b) and intensity-related effects (Figure 5b/c) of the US. All locations are reported using Montreal Neurological Institute (MNI) coordinates (XYZ_MNI_). As expected, heat stimulation was followed by larger activation in widespread insular and opercular areas, with the highest peak in the dorsal posterior insula (XYZ_MNI_ 35.5/-17.9/21.4, T = 12.2, p[corr.] = ∼0). Notably, a conjunction of both heat and sound main effects shows activation in the central operculum (XYZ_MNI_ 53.0/-10.3/15.1, T = 8.3, p[corr.] = 8 x 10^-13^), dorsal anterior insula (XYZ_MNI_ 37.6/18.4/-7.0, T = 5.6, p[corr.] = 2 x 10^-05^), and several regions in between peaks for both modalities.

**Figure 5.**
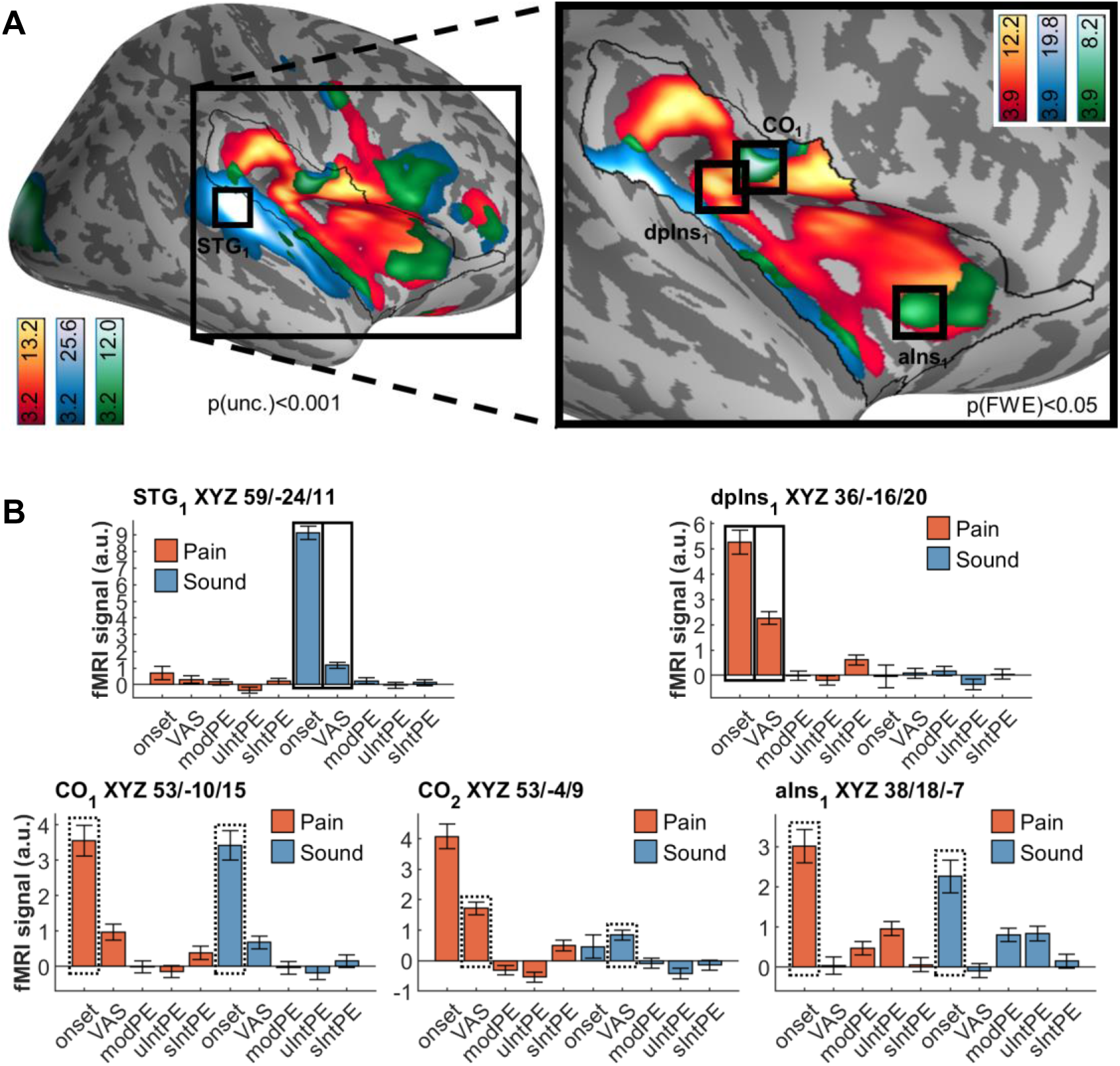
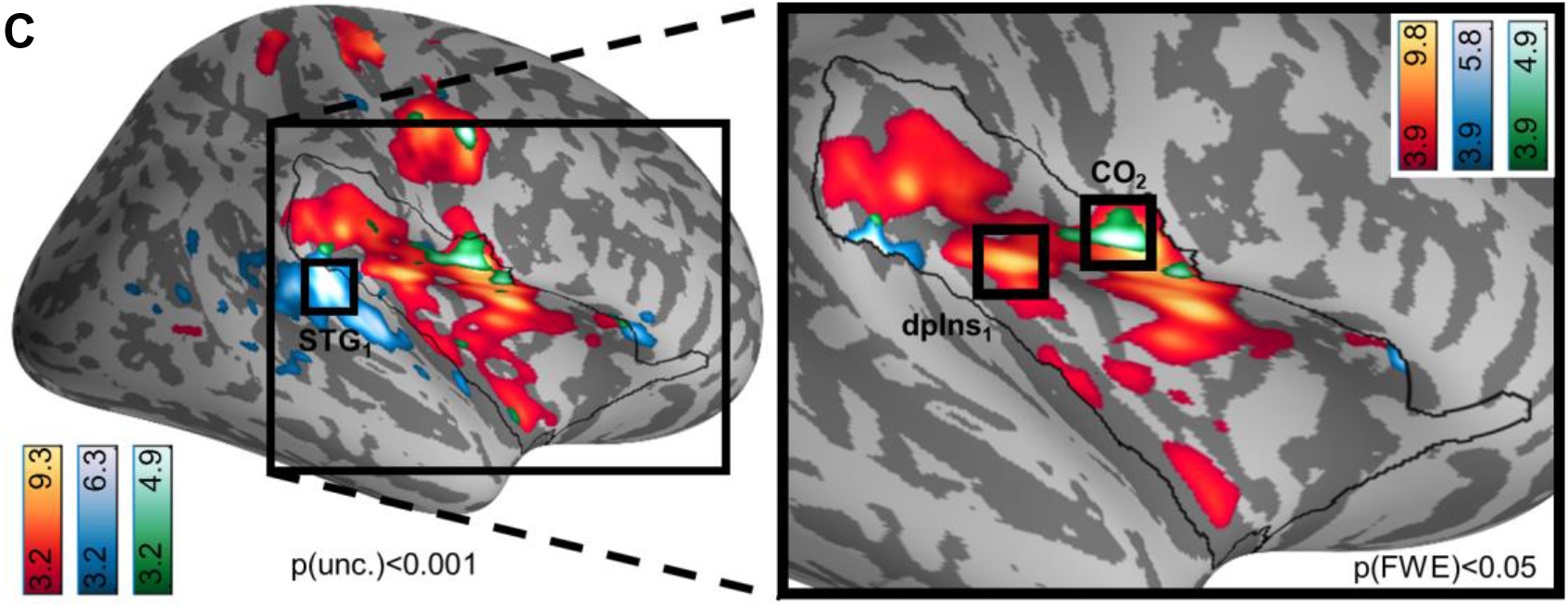
Brain activation following pain (red/yellow) and sound (blue), including overlaps as per conjunction analyses (green). Activations are overlaid on an average brain surface; for display purposes, activations in the whole brain lateral view are thresholded at p[uncorr.] < 0.001. The black line in the zoomed-in view delineates the region of interest and includes activations within the small volume FWE-corrected at p[corr.] < 0.05. Peaks are shown for small volume only; bar plots show beta weights of BOLD activation obtained from a general linear model (see Materials and Methods) from the respective peaks See supporting information for peak positions in whole brain (Supporting Figure 4, Supporting Figure 6), and brain *volume* slices (Supporting Figure 5, Supporting Figure 7). **(A)** Differential and shared activation following painful heat stimulation and loud sound stimulation. Peak activation following heat is located in (peri)insular areas contralateral to stimulation, namely the dorsal posterior insula (dpIns1), and extending through the central and parietal opercula. Peak activation following sound is located in the superior temporal gyrus. Common activation (green) is located in the central operculum (CO1) and dorsal anterior insula (aIns1), among other regions. **(B)** fMRI signal (arbitrary units) for peaks detected in panel a (US onset effects) or c (parametric modulation by ratings). **(C)** Differential and shared correlations with pain ratings (for heat) and unpleasantness ratings (for sound). Activation correlated with pain ratings is focused on the dorsal posterior insula (dpIns1). Activation correlated with sound ratings is focused on the superior temporal gyrus. Conjunction activation peaks in central operculum (CO2) and precentral gyrus. fMRI signal regressor labels: VAS, visual analogue scale; PE, prediction error; modPE, modality PE; uIntPE, unsigned intensity PE; sIntPE, signed intensity PE.

Next, we tested for fMRI responses correlated with stimulus perception, i.e. pain and sound VAS ratings (Figure 5b/c). For pain ratings, associations arose in the dorsal posterior insula (XYZ_MNI_ = 35.2, y = -17.4, z = 18.6, T = 7.2, p[corr.] = 1 x 10^-09^). For sound ratings, we observed a peak directly adjacent to the small surface (XYZ_MNI_ 59.8, y = -33.9, z = 5.4, T = 4.8, p[corr.] = 0.016). Common activation between pain and sound ratings peaked in the central operculum (XYZ_MNI_ 53.2, y = - 2.7, z = 8.9, T = 4.8, p[corr.] = 0.001). Of note, the central operculum peak (CO_2_ in Figure 5c) is located slightly anterior to that found for the modality conjunction (CO_1_ in Figure 5a) but shows barely any sound modality activation; conversely, peak aIns1 indicates that no intensity effects are encoded here. See supporting information for additional activations (Supporting Figure 4, Supporting Figure 6).

### Unsigned intensity prediction errors

Having ascertained strictly stimulus-related effects, our next analysis included an investigation of *unsigned* intensity PEs within and between either modality (Figure 6). The guiding question here was whether any differences and commonalities between the modalities would emerge. Since we used the actual expectation queried from subjects, “prediction error” here means that subjects explicitly expected one intensity but received the other. Consequently, the unsigned PE implies some extent of surprise.

**Figure 6.**
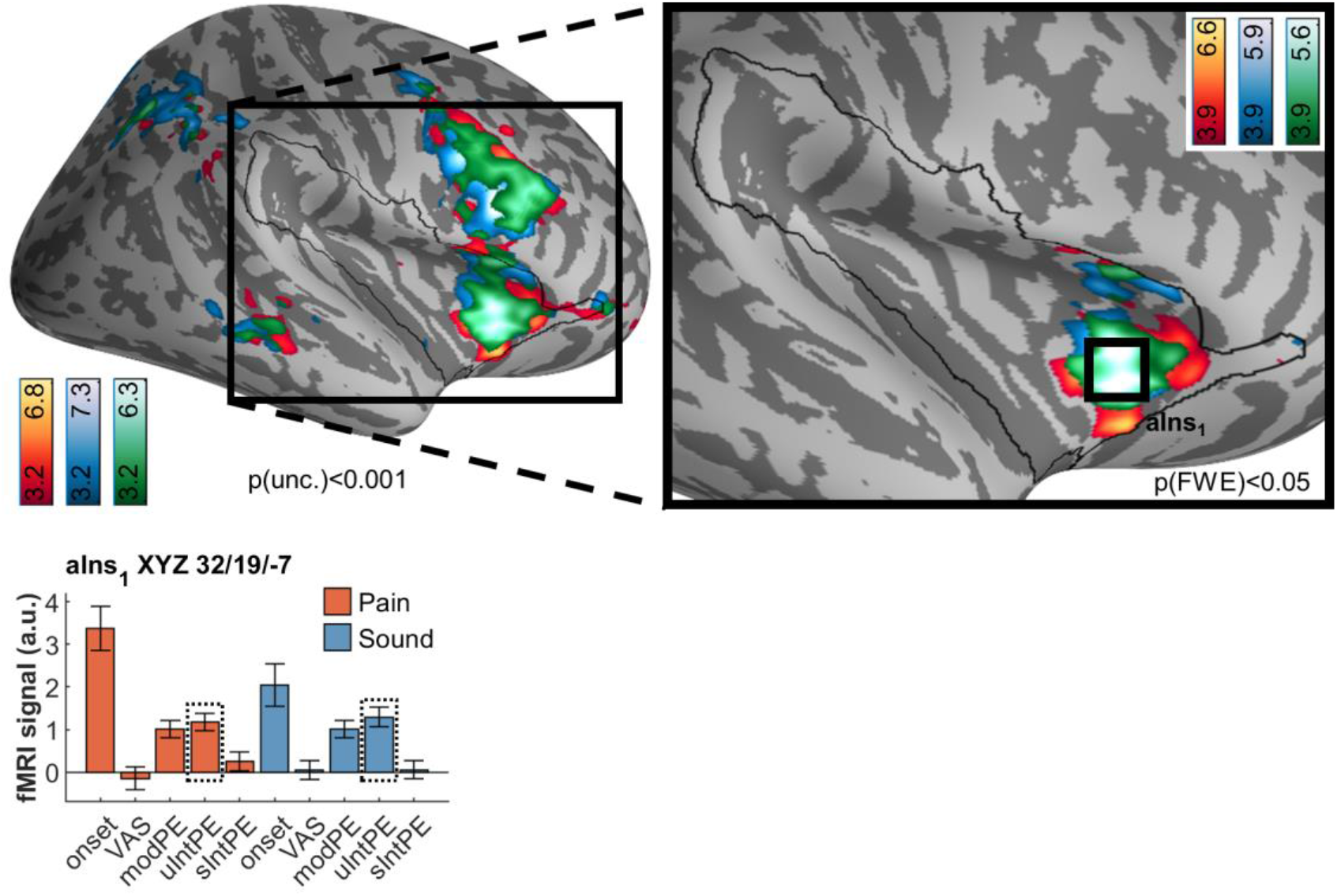
Brain activation following unsigned intensity prediction errors in pain (red/yellow) and sound (blue), including overlaps as per conjunction analyses (green). Peak activation following either modality is located in the anterior insula (aIns1) and is subsumed in the common activation. Activations are overlaid on an average brain surface; for display purposes, activations in the whole brain lateral view are thresholded at p[uncorr.] < 0.001. The black line in the zoomed-in view delineates the region of interest and includes activations within the small volume FWE-corrected at p[corr.] < 0.05. See supporting information for peak positions in whole brain (Supporting Figure 8), and brain *volume* slices (Supporting Figure 9). fMRI signal regressor labels: VAS, visual analogue scale; PE, prediction error; modPE, modality PE; uIntPE, unsigned intensity PE; sIntPE, signed intensity PE.

In both modalities, widespread activation was observed. However, conjunction analyses revealed that the majority of the observed activation actually overlapped between the modalities (green in Figure 6). The anterior insula constituted the dominant cluster of this overlap, with symmetric bilateral peaks (XYZ_MNI_ = 34.6/23.5/-1.5, T = 5.8, p[corr. wb.] = 1 x 10^-04^); whole brain-significant frontal (medial and lateral), temporal and parietal activation was also observed (Supporting Figure 8).

Two aspects were of particular interest to us considering unsigned intensity PE results: First, that brain activation related to unsigned intensity PEs (Figure 6) was distinct from the intensity-related activation (Figure 5). Second, the fMRI signal of the common activation in the anterior insula clearly indicated that modality PEs are likewise encoded in this area.

### Modality prediction errors

Following these two observations, we proceeded to investigate the nature of the overlap between the two types of PE. Like with unsigned intensity PEs, we observed widespread activation following each modality PE separately (Figure 7). Likewise, all unimodal activation is subsumed in the conjunction analysis, which indicates a large dorsal anterior insula cluster in our region of interest (XYZ_MNI_ 32.3/22.4/-3.4, T = 5.4, p[corr.] = 5 x 10^-05^). Beyond this region, widespread common activation is observed, for example, in the superior parietal lobule, precuneus, temporo-parietal junction, middle frontal gyrus and frontal operculum, and medial orbital gyrus (Supporting Figure 10).

**Figure 7.**
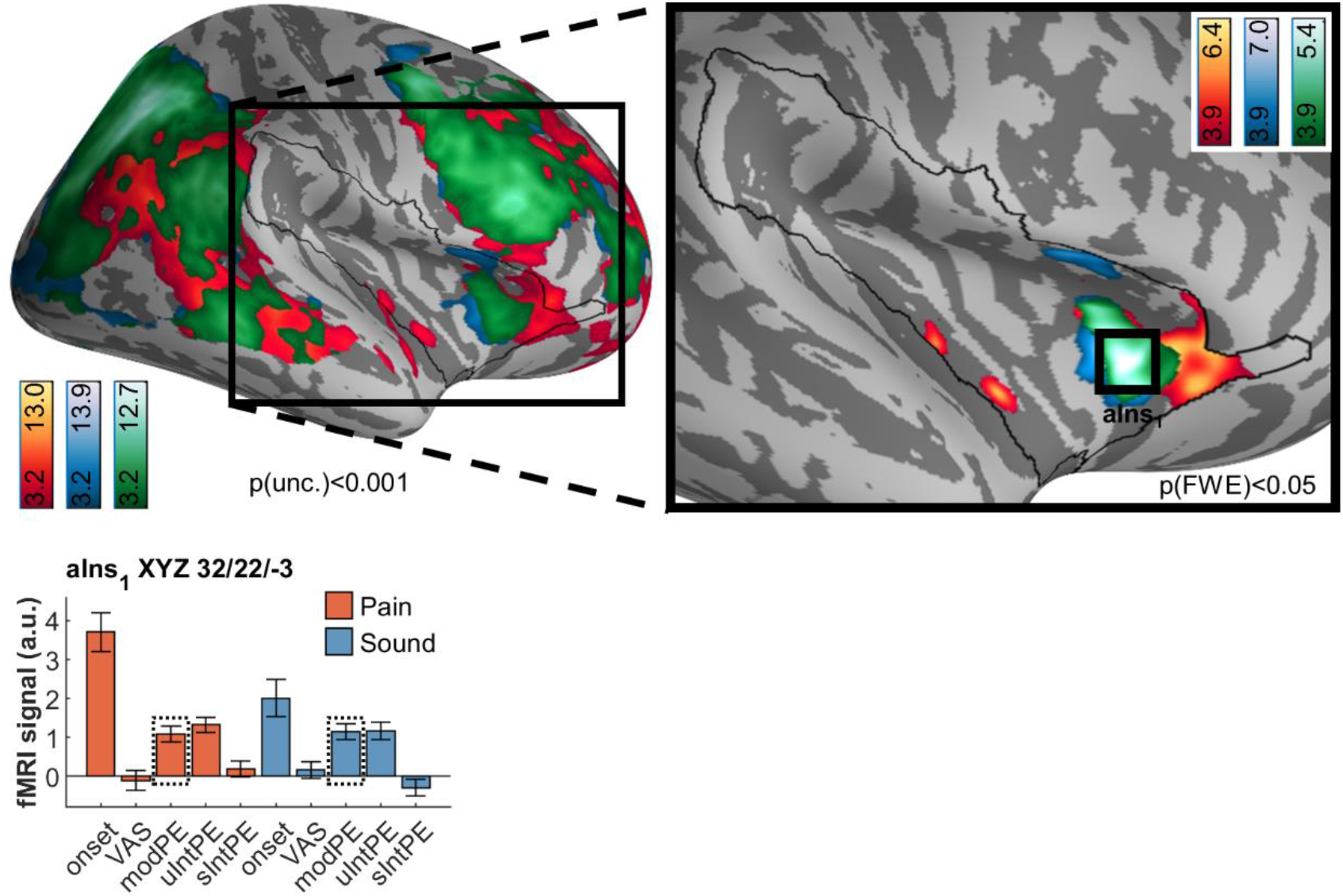
Brain activation following modality prediction errors in pain (red/yellow) and sound (blue) activation, including overlaps as per conjunction analyses (green). As with unsigned intensity PEs, peak activation following modality PEs in either modality is located in the anterior insula (aIns1) and is largely subsumed in the common activation. Activations are overlaid on an average brain surface; for display purposes, activations in the whole brain lateral view are thresholded at p[uncorr.] < 0.001. The black line in the zoomed-in view delineates the region of interest and includes activations within the small volume FWE-corrected at p[corr.] < 0.05. Peaks are shown for the small volume only. See supporting information for peak positions in whole brain (Supporting Figure 10), and brain *volume* slices (Supporting Figure 11). fMRI signal regressor labels: VAS, visual analogue scale; PE, prediction error; modPE, modality PE; uIntPE, unsigned intensity PE; sIntPE, signed intensity PE.

### Overlap of unsigned prediction errors

As a next step, we wanted to more formally assess the apparent overlap between both types of unsigned PEs. To do so, we simply computed the conjunction between unsigned intensity and modality PE (Figure 8). This analysis corroborated the anterior insula peak determined by separate analyses above. Furthermore, activation extended dorsally through the middle frontal gyrus and also included medial prefrontal areas adjacent to the dorsal anterior cingulate cortex.

**Figure 8.**
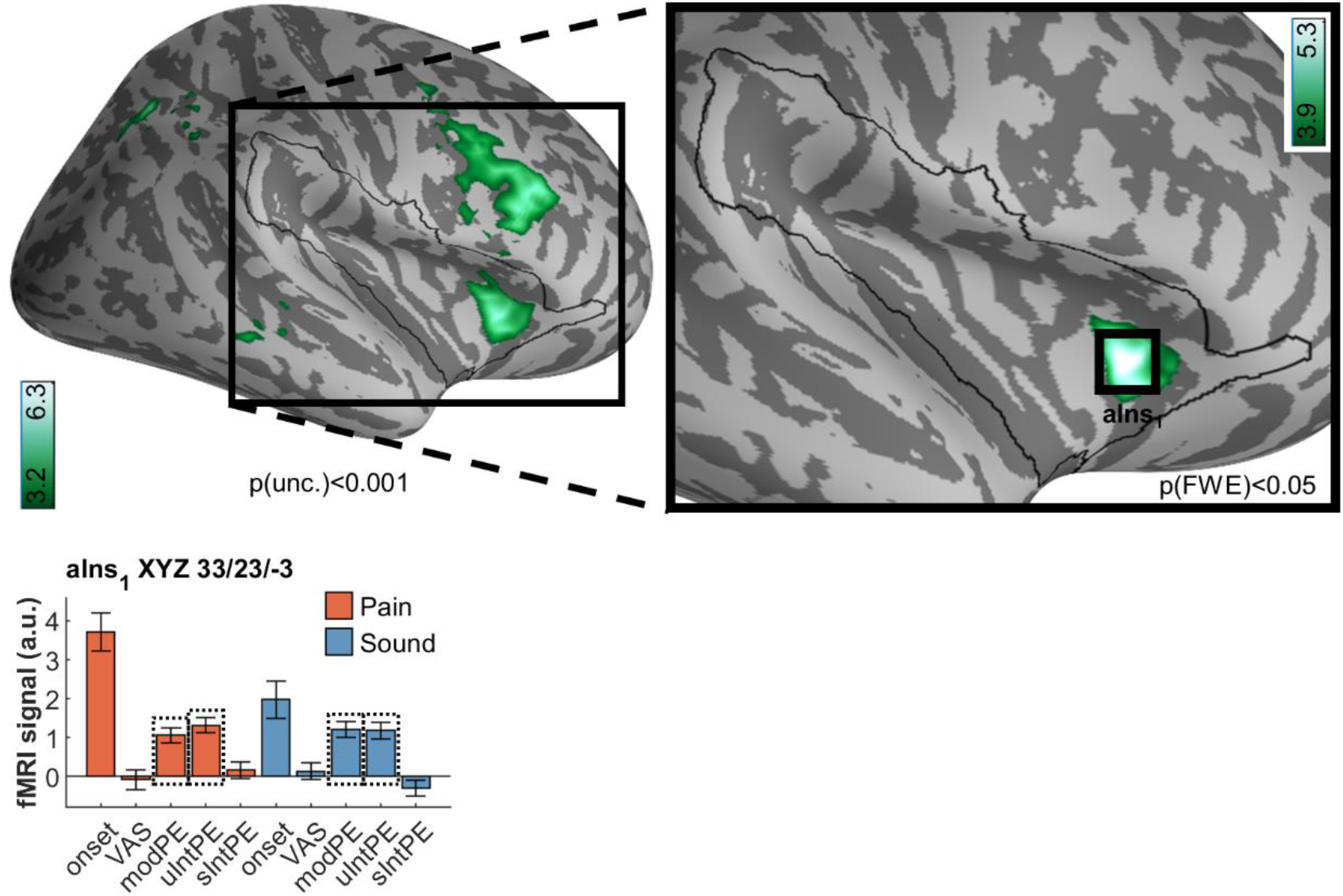
Common brain activation associated with unsigned intensity and modality prediction errors. The fMRI signal plot shows that the peak in the anterior insula (aIns1) encodes PEs for every contrast included in the conjunction. Activations are overlaid on an average brain surface; for display purposes, activations in the whole brain lateral view are thresholded at p[uncorr.] < 0.001. The black line in the zoomed-in view delineates the region of interest and includes activations within the small volume FWE-corrected at p[corr.] < 0.05. fMRI signal regressor labels: VAS, visual analogue scale; PE, prediction error; modPE, modality PE; uIntPE, unsigned intensity PE; sIntPE, signed intensity PE.

### Signed intensity prediction errors

After ascertaining the effects for *unsigned* PEs for both intensity and modality, the final question for our fMRI data referred to differences and commonalities following *signed* intensity PEs, i.e. correlations of brain activation with higher-than-expected intensity (Figure 9). For pain, we observed an activation in the dorsal posterior insula (XYZ_MNI_ 36.4/- 17.3/15.8, T=4.0, p[corr.]=0.023). The dorsal posterior insula is an area considered of fundamental importance for the processing of pain intensity [24,28,31]. For sound itself, the peak activation was observed outside the region of interest, in the middle temporal gyrus (XYZ_MNI_ 49.4/-16.6/-13.4, T=4.1, p[uncorr.]=2 x 10^-05^) (see Figure 6). Within the region of interest, sound-related activation was found in the anterior insula (XYZ_MNI_ 36.7/11.0/-10.2, T=4.2, p[corr.]=0.015). Notably, these are adjacent to the unsigned PE activations (Figure 6 through Figure 8). All signed intensity PE peaks, both for pain and sound, show no significant representation of a signed PE in the other modality (see opposite sIntPE fMRI signals in Figure 9). Consequently, a conjunction analyses revealed no overlap.

**Figure 9.**
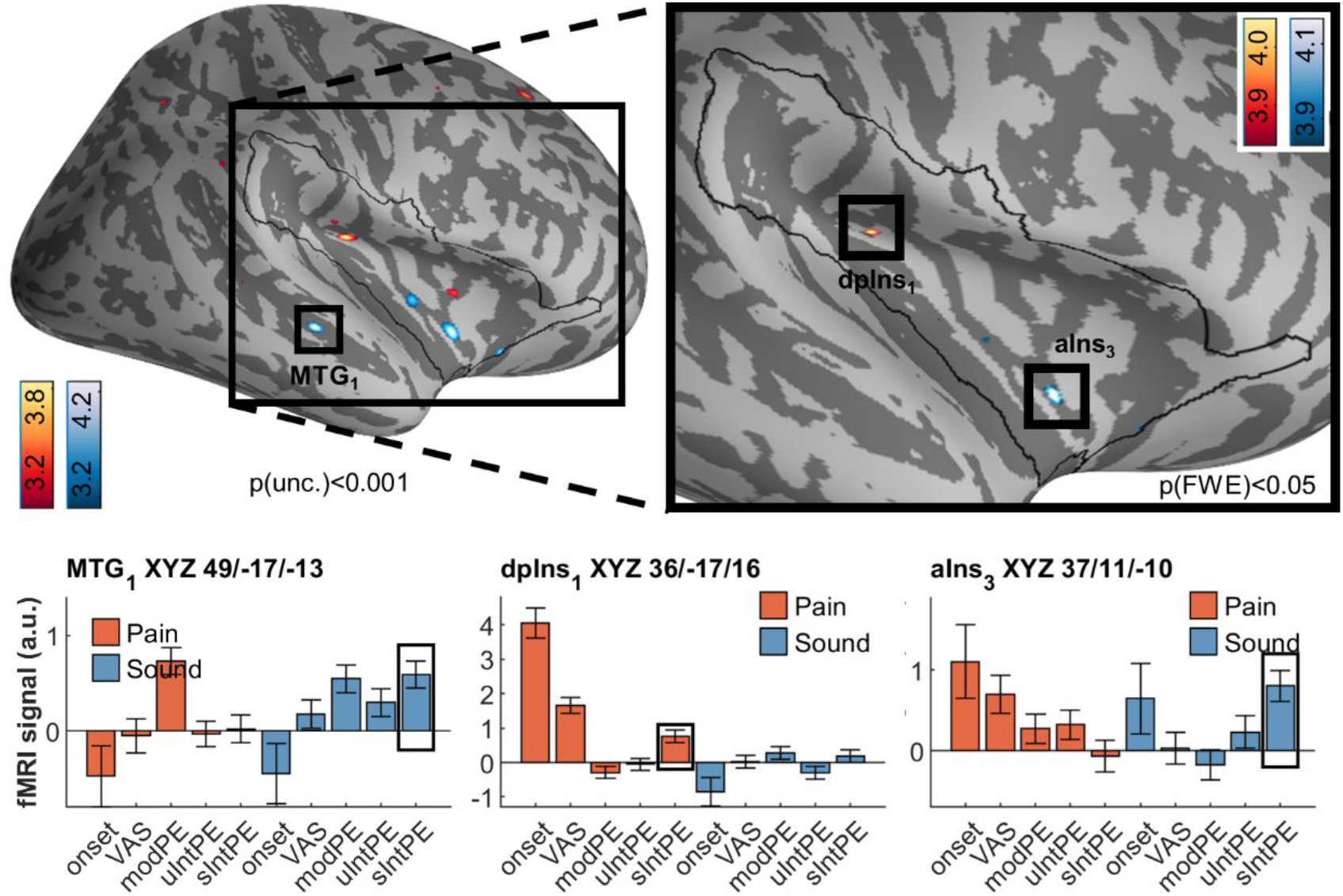
Brain activation associated with by signed intensity prediction errors in pain (red/yellow) and sound (blue), including overlaps as per conjunction analyses (green). Contrary to uIntPEs, circumscribed activation was detected for pain sIntPEs without any overlap with sound sIntPEs. Peak activation is located in the dorsal posterior insula (dpIns1). For sound, several clusters in the anterior insula (e.g. aIns3) were found, as well as middle temporal gyrus (MTG1). Activations are overlaid on an average brain surface; for display purposes, activations in the whole brain lateral view are thresholded at p[uncorr.] < 0.001. The black line in the zoomed-in view delineates the region of interest and includes activations within the small volume FWE-corrected at p[corr.] < 0.05. fMRI signal regressor labels: VAS, visual analogue scale; PE, prediction error; modPE, modality PE; uIntPE, unsigned intensity PE; sIntPE, signed intensity PE.

In summary, the unsigned intensity PEs for pain and sound, as well as their modality PEs, strongly overlap in the anterior insula (Figure 6), whereas signed intensity PEs are accompanied by pain-dedicated activation in the dorsal posterior insula (Figure 9).

## Discussion

Using a Pavlovian learning paradigm with frequent reversals within and across aversive modalities in combination with SCR recordings and high resolution fMRI, we were able to investigate signed and unsigned representations of PEs in the human brain. The data showed an unsigned representation of intensity PEs in the anterior insula indistinguishable for pain and aversive sounds, supporting a role of the anterior insula in coding unspecific arousal or salience. In addition, the same part of the anterior insula also strongly activated for PEs concerning stimulus modality. Most importantly, we could identify a circumscribed part of the dorsal posterior insula representing a signed PE for pain only, collocated with areas processing pain intensity per se.

The parallel assessment of SCR, behavioral ratings for both expectation and outcome, as well as fMRI recordings allowed us to investigate PEs in a multimodal fashion. Previous studies investigated PEs using cue-based pain paradigms [12,19,21,32]. In these paradigms, a cue predicts a pain intensity with a certain probability. However, the probability also determines the number of trials in which a PE occurs. This can lead to unbalanced designs in which certain PEs occur much more frequently than others. In addition, the fixed association of a specific cue with an outcome risks that specific features of the cue influence PE processing. Adopting a Pavlovian transreinforcer paradigm ameliorates these shortcomings, and requires frequent relearning of contingencies and thus generates frequent PEs [22, 23]. By defining a Markovian transition structure, we also controlled the nature of reversals; we confined our experiment to within-intensity/between-modality, and between-intensity/within-modality reversals. Finally, introducing two CS in our task increased task difficulty.

We explicitly included expectation ratings, which allowed us to use the difference between the US and its expectation as a rating-derived PE [22]. Compared to model-derived PEs, this can account for within-subject differences in learning and can also capture PEs in erratic behaviors difficult to model in formal reinforcement learning models.

Although we aimed to perfectly match salience between stimulus modalities, high intensity painful stimuli lead to higher SCR activation compared to low pain or either sound intensity (Figure 4), even though average SCR amplitudes between modalities were not statistically different. Technically, this is related to the fact that we were not able to increase sound pressure levels above a certain level [33] to avoid harm for the volunteers. However, the fMRI signal changes in the anterior insula for unsigned intensity PEs were similar for pain and sound, suggesting that the residual differences in SCR did not affect our results (Figure 6, Figure 7, Figure 8). In addition, previous accounts [34] have indicated that higher salience enhances memory performance. We tested this and observe no such effect: learning performance did not substantially differ between any of the US groups (Supporting Figure 14).

We have replicated findings concerning pain-related activation in the dorsal posterior insula/parietal operculum and sound-related activation in the superior temporal gyrus [24]. Previously, these areas showed a clear effect of pain and sound stimulation, respectively, but a crucial intensity-related increase in activation that is shallower or absent in non-noxious intensities. In contrast to the previous study, we see a stronger correlation of the BOLD response to sound ratings, possibly owing to the higher intensities employed here.

Also in agreement with previous studies, we observed an unsigned intensity PE for pain in the anterior insula [12,19,21]. The novel contribution is the fact that stimuli in different modalities (i.e. pain and aversive sounds) [24] lead to the same activations in the anterior insula, with similar magnitudes. To our surprise, strong activation in the anterior insula was also observed for modality PEs (expect pain and receive sound, and vice versa). fMRI signals for unsigned intensity PEs and modality PEs were very similar in magnitude. This disconfirms our hypothesis that at the level of the insula, modality PE carries less difference in salience between the expected and the real outcome, as compared to an unsigned intensity PE. Rather, it seems that surprise from unexpected sensory modalities is as much a source of anterior insula activation as from unexpected intensities. Our findings suggest that modality and unsigned intensity PEs are largely modality-neutral, and support findings that the anterior insula is richly interconnected part of the salience and attentional network involved in decision-marking, error recognition and generally the guidance of flexible behavior [35–39]. Indeed, the large-scale activation following modality PEs and unsigned intensity PEs themselves does not correspond to any single network description, but seems to involve all of the above; possibly, different dynamics are at play over the course of the stimulation, which do not allow for the disentangling of single networks. In fact, recent meta-analytic evidence of resting-state functional connectivity points to the existence of a pain-related network centered on the anterior insula [40]. The activation associated with both pain-related (posterior insula) activation, and that associated with PE-related (anterior insula) activation correspond well with connectivity gradients observed along the posterior-anterior axis [41–43].

It is known that SCR predominantly shows arousal and similar effects, but is relatively insensitive concerning valence [25–27,44,45]. Here, SCR following unsigned or signed intensity PEs was little different from SCR following no PEs, while SCR following modality PEs was much higher. This might indicate that modality PEs provide a highly salient a teaching signal even in the absence of intensity differences (Supporting Figure 3).

A signed representation of an intensity PE for pain is a crucial teaching signal in reinforcement learning, as it is important to dissociate a low threat from a high threat stimulus. Such a representation for pain could plausibly be located in an area adjacent the anterior insula part representing unsigned intensity PEs and modality PEs. Alternatively, this representation could be located closer to representations of pain intensity: Coding of signed intensity PEs within areas coding for stimulus intensity per se was observed using a similar Pavlovian transreinforcer paradigm in the olfactory domain [23]. Indeed, our data show that a signed intensity PE for pain is represented in a part of the dorsal posterior insula [24, 28]. Interestingly, we also identified a similar representation of a signed intensity PE for aversive sounds in or adjacent to primary auditory cortices [46, 47], namely the middle temporal gyrus and temporal operculum. It also seems indicative of the more general involvement of the insula in pain perception [48] that the signed intensity PE in pain has little to none sound-related activation at all, whereas the signed intensity PE in sound includes some pain intensity-related activation.

At most, the clear spatial dissociation of intensity PEs for pain and sounds furthermore indicates a specificity of the signal; at least, it stands in marked contrast with the large overlap of activation for unsigned intensity and modality PEs in the anterior insula. Powerful learning models can utilize both a signed PE to update their predictions and an unsigned PE to update their learning rate [10,17,18]. Our results provide a neuronal basis for these models as we were able to reveal the simultaneous representation of both a signed and unsigned PE signal in spatially distinct regions of the insula.

Due to the task-inherent structure, signed pain intensity PEs can be correlated with actual pain intensity [49]. This collinearity can be remedied by orthogonalizing regressors in the general linear model used for fMRI analysis. However, this arbitrarily assigns the shared variance to either of the two correlated regressors, depending on the order of the serial orthogonalization [50]. Therefore, we refrained from any orthogonalization in our analysis and thus only reveal areas that show unique variance tied to the regressors, including the signed intensity PEs for pain.

In conclusion, our data provides clear evidence of anterior insula-centered, modality-independent *unsigned* PEs, not only concerning mismatched stimulus intensities across modalities, but also across sensory modalities themselves. Equally important, *signed* intensity PEs were associated with activation in or adjacent to sensory areas highly dedicated to unimodal processing. Neuronal data from both sources are the basis for reinforcement learning and further enhance our understanding of the functional synergies within the insula. Importantly, pathological learning mechanisms [1, 9] and abnormalities in anterior insula-related function have been reported in chronic pain [40, 51]. Our data therefore offers the possibility that a misrepresentation of PEs constitutes a potential mechanism in pain persistence.

## Materials and Methods

The protocol conformed to the standards laid out by the World Medical Association in the Declaration of Helsinki and was approved by the local Ethics Committee (Ethikkommission der Ärztekammer Hamburg, vote PV4745). Participants gave written informed consent prior to participation and were aware of all aspects of the protocol except the randomized time point of reversal trials.

### Subjects

Forty-nine healthy volunteers (Sex 27f:22m, Age 26.2±4.5) were recruited through online advertisements (www.stellenwerk.de) and word of mouth. They were screened concerning study- and MR-specific exclusion criteria as follows:

- Age younger than 18, older than 40
- Insufficient visual acuity (correction with contact lenses only)
- Conditions disqualifying for MR-scanners (e.g. claustrophobia, wearing a pacemaker)
- Ongoing participation in pharmacological studies, or regular medication intake (e.g. analgesics)
- Analgesics use 24h prior to the experiment
- Pregnancy or breastfeeding
- Chronic pain condition
- Manifest depression (as per Beck Depression Inventory II, cutoff 14 [52])
- Somatic symptom disorder (as per Patient Health Questionnaire, cutoff 10 [53])
- Other neurological, psychiatric or dermatological conditions
- Inner ear conditions
- Head circumference >60 cm (due to MR scanner coil/headphone constraints)

Eligible subjects were scheduled for a single lab visit. Experiments were conducted from October 2019 through March 2020. Statistics characterizing the sample are listed in Supporting Table *2*.

### Overview of the experiment

The sequence of measurements and timings of the protocol are displayed in Figure 1, while aspect pertaining to CS characteristics as well as contingencies are displayed in Figure 2. The experiment lasted about 2.5 h. The experiment followed a full cross-over design, with every subject participating in all conditions. Subjects learned associations of conditioned stimuli (CS) and unconditioned stimuli (US; painful heat or loud sound). These associations eventually changed in an unforeseeable manner and then had to be relearned. The experiment was run in a single visit, but split into two sessions to reduce subject fatigue and carry-over effects. Prior to the experimental sessions, subjects were calibrated according to their pain and sound sensitivity. At the start and the end of the experiment, subjects filled out psychological questionnaires outside the scanner. Electrodermal activity was measured throughout the experimental sessions.

### Unconditioned stimuli

Heat stimuli were delivered using a CHEPS thermode (Medoc, Ramat-Yishai, Israel) attached to the volar forearm. Basic stimulus parameters included a 32°C baseline temperature and 10°C/s rise and fall rates. Sound stimuli were delivered using MR-compatible headphones (MR confon, Magdeburg, Germany). A pure sound (frequency 1000 Hz, sampling rate 22050 Hz) was generated during runtime using MATLAB.

### Calibration of unconditioned stimulus intensities

Prior to the experiment proper, subjects underwent US calibration to determine two intensities at VAS 25 and VAS 75 for both modalities (heat and sound). During the experiment, only these four stimuli were used. All stimuli lasted 3s at plateau, except for four 10s long, low-intensity preexposure stimuli used for familiarization and pre-heating of the skin.

Heat and sound stimuli were presented and rated in an analogous fashion. Like in a previous study comparing neuronal responses to the two modalities [24], we used the descriptor “painfulness” for heat, while we used the descriptor “unpleasantness” for sound. After calibration, all stimuli were above the respective pain and unpleasantness thresholds and were therefore displayed on simple 0 to 100 visual analogue scales (VAS) for both modalities.

For heat, anchors were displayed for “minimal pain” (0) and “unbearable pain” (100). Pain was defined as the presence of sensations other than pure heat intensity, such as stinging or burning [54].

For sound, subjects were instructed to rate between anchors labelled “minimally unpleasant” (0) and “extremely unpleasant” (100). Unpleasantness was defined as a bothersome quality of the sound emerging at a certain loudness.

During the calibration procedure performed in the running MR scanner, two stimulus intensities each were obtained for the heat and sound modality (low/high pain and low/high noise). Heat stimuli ranged from 43 to 49°C, sound stimuli ranged from 89.1 through 103.0 dBA. Calibration was constrained such that subjects had to reach a certain

- minimum physical intensity (43°C for heat, 20% system volume for sound, n=1 received 10%)
- minimum physical difference between the VAS 25 and 75 stimuli (1.5°C for heat, 15% system volume for sound; n=1 received 1°C, n=8 received 10%)

If either condition was not met, physical intensities were automatically adjusted to the minimum (e.g., if subject reported VAS 25 for 41°C, temperature was raised to 43°C). Furthermore, to ensure discriminability within stimulus modalities, subjects had the calibrated US played back to them and were explicitly asked three questions, namely that both intensities of the respective modality

- were painful (for heat) or unpleasant (for sound)
- were perspectively tolerable throughout repeated trials in two sessions
- were easily discriminable.

If either question was answered in the negative, the calibrated intensities were adjusted, but never below the minimum requirements listed above.

### Learning protocol

Learning the CS-US associations was designed as a Pavlovian transreinforcer reversal learning task [22, 23]. Two CS would independently predict one of four US, namely two intensities of painful heat and two intensities of unpleasant sound. Subjects were presented with one of the two CS (Figure 2c and d) and then asked to choose which of the four US they believed to be preceded by it (symbols in Figure 2b). After making their choice, they would actually be exposed to one of the four US (see Figure 1c for trial structure). If they were correct, no further learning was required; if not, they would have the opportunity to learn the correct association for the next occurrence of the CS. They would then rate their pain or unpleasantness on a 0-100 visual analogue scale (VAS), as during US calibration. Both CS signified an independent sequence of associations with the US. Both CS were randomly drawn for each subject from a library of eight fractal pictures (Figure 2a). Which of the two CS was presented in each trial was fully randomized, as were the US for the respective initial associations, and the display order of the US prediction rating.

Crucially, after a number of trials with deterministic CS-US association, the association underwent an unannounced reversal either in terms of intensity (previously low US intensity would now be high, or vice versa), or modality (previous pain US would now be a sound US, or vice versa) (Figure 2c and d). The number of trials that an association was upheld was randomly determined from [3, 3, 4, 5] (i.e. 3.75 trials on average). After each reversal, subjects therefore made an error in predicting the following US, and subsequently had to learn the new association. As reversals on both dimensions were precluded, each session included eight reversals per CS to cover all possible reversals. Task performance was assessed by the percentage of correct predictions.

### Psychological questionnaires

Prior to and immediately after the experiment, subjects filled out several questionnaires assessing state and trait psychological constructs. These are listed in Supporting Table *2* alongside statistics characterizing the sample.

### Psychophysiological recordings

Electrodermal activity was measured with MRI-compatible electrodes on the side of the left hand opposite the thumb. Electrodes were connected to Lead108 carbon leads (BIOPAC Systems, Goleta, CA, USA). The signal was amplified with an MP150 analog amplifier (also BIOPAC Systems). It was sampled at 1000 Hz using a CED 1401 analog-digital converter (Cambridge Electronic Design, Cambridge, UK) and downsampled to 100 Hz for analysis.

Analysis was performed using the Ledalab toolbox for MATLAB [55]. Single subject data were screened for artifacts which were removed if possible by using built-in artifact correction algorithms. Of 47 subjects, 1 was excluded due to equipment malfunction, 9 due to skin conductance non-responsiveness. From the remaining 37 subjects, a total of 101 of 6016 segments (1.7%) were excluded due to unsalvageable artefacts. Using a deconvolution procedure, we computed the driver of phasic skin conductance (skin conductance responses, SCR). Stimulus phase response windows were offset between the two stimulus modalities [24] – we attribute an earlier onset following acoustic stimulation to reduced latency from the delivery system and neuronal transmission. To determine response windows, we obtained the times for average peaks of the respective modality, and selected the data range ±1.25 s: For pain, response windows were set between 2.42 s and 4.92 s, and between 1.15 s and 3.65 s for sound. SCR segments were log- and z-transformed within subjects to reduce the impact of intra- and interindividual outliers [25]. Subsequently, segments were averaged within subjects for several conditions corresponding to the behavioral performance of subjects (e.g. intensity PE following low painful stimulation, or high painful stimulation). SCR was used because it is an objective measure of general sympathetic activity, and therefore a measure of arousal, stimulus salience and several associated psychological processes [25,26,45,56,57]. It is routinely used in assessing painful [12,24,58] as well as acoustic stimulation [59].

### fMRI acquisition and preprocessing

Functional and anatomical imaging was performed using a PRISMA 3T MR Scanner (Siemens, Erlangen, Germany) with a 20-channel head coil. An fMRI sequence of 56 transversal slices of 1.5 mm thickness was acquired using T2*-weighted gradient echo-planar imaging (EPI; 2001 ms TR, 30 ms TE, 75° flip angle, 1.5×1.5×1.5 mm voxel size, 1 mm gap, 225×225×84 mm field of view, simultaneous multislice imaging with a multiband factor of 2, and an acceleration factor of 2 with generalized autocalibrating partially parallel acquisitions reconstruction). Additionally, a T1-weighted MPRAGE anatomical image was obtained for the entire head (voxel size 1×1×1 mm, 240 slices).

For each subject, fMRI volumes were realigned to the mean image in a two-pass procedure, and non-linearly co-registered to the anatomical image using the CAT12 toolbox for SPM (Christian Gaser & Robert Dahnke, http://www.neuro.uni-jena.de/cat/). In short, this novel non-linear coregistration segments both the mean EPI and the T1 weighted image and performs a nonlinear spatial normalization of the segmented tissue classes from the mean EPI using the segmented tissue classes from the T1 scan as a template. Finally, individual brain surfaces were generated, using CAT12.

### General statistical approach

Unless otherwise noted, analyses except the fMRI analyses were performed using linear mixed models with random intercept using trial-by-trial parameters. In the case of mixed (within/between) descriptive statistics, standard errors were calculated using the Cousineau-Morey approach [60]. The significance level for analyses of behavioral and psychophysiological data was set to p = 0.05.

### Analysis of imaging data

Subject-level analyses were performed on the 3D (volume) data in native space without smoothing, as required for surface mapping. We computed a general linear model with a canonical response function to identify brain structures involved in the processing of each stimulus modality, and corresponding to various predictions and PEs inherent in the protocol. Realignment (motion) parameters were included as nuisance variables, to further mitigate motion-related artifacts.

A general linear model was set up with one regressor for stimulus main effects in each modality (heat or sound), and a parametric modulator each for pain or unpleasantness (using behavioral ratings). An additional three parametric modulators for each modality were entered for modality PEs and intensity PEs: Modality PEs were entered unsigned due to their non-parametric nature, whereas intensity PEs were entered both unsigned (absolute) and signed. All parametric modulators were z-scored within subjects and sessions. In either model, global or sequential orthogonalization between regressors were turned off to preserve only the unique (non-shared) variance components [23, 50]. This approach allows for the interpretation of consecutively entered parametric modulators even if correlations to previous regressors exist.

We opted for surface-based analyses of fMRI data to enhance discrimination between modalities processed in adjacent brain regions [24]; for an example of pseudo-overlap detected across the sylvian fissure, see Supporting Figure 5 (row 3), particularly in slices -28 through -16. Results from subject-level analyses were mapped to brain surfaces obtained via the CAT12 segmentation procedure. The mapped subject-level results were then resampled to correspond to cortical surface templates, and smoothed with a 6 mm full width-half maximum 2D kernel. Group-level within-subjects analyses of variance were performed including the mapped contrasts. The original, unmapped contrasts were used for volume-based group-level analyses to assess subcortical activation. Volume results were then warped using DARTEL normalization and smoothed with a 6 mm full width-half maximum 3D kernel. Volume-based results are provided in the supporting information and referenced where relevant.

Contrasts employed for any of the analyses were either performed against low-level baseline (e.g. Pain>0), as a conjunction of a differential modality contrast and one against low-level baseline (e.g. Pain>Sound ∧ Pain>0), or as a conjunction of both modalities (e.g. Pain ∧ Sound).

### Regions of interest and statistical correction of imaging results

As laid out above and because pain is the modality of interest in this study, we focused the analyses on the contralateral (right) periinsular cortices as regions of interest used for small volume correction of significance level [12,19,24]. The region of interest included the entire insular cortex (dorsal hypergranular, dorsal granular, dorsal dysgranular, dorsal agranular ventral dysgranular/granular, ventral agranular), as well as dorsally adjacent areas of the parietal operculum (A40rv), central operculum (A1/2/3ll, A4tl) and frontal operculum (A44op, A12/47l). It was created using the Human Brainnetome Atlas [61]. Results were considered after correction for family-wise error rate of p < 0.05 within the region of interest (denoted p[corr.]), or after correction for whole brain/all vertices (denoted p[corr. wb.]), unless otherwise noted.

## Acknowledgements

This work was supported by ERC-AdG-883892-PainPersist and DFG SFB 289 Project A02 (Project-ID 422744262–TRR 289). We thank Thorsten Kahnt for comments and scripts concerning the randomization procedure, Saša Redžepović for providing scripts used for CS fractal generation, Jürgen Finsterbusch, Katrin Bergholz, Waldemar Schwarz and Kathrin Wendt for technical assistance during MR data collection, and Alina Schaefer and Jannis Petalas for their assistance with data collection.

## Author contributions

B.H. and C.B. conceived and designed the study, analyzed and interpreted the data, and wrote the manuscript. B.H. performed the experiments.

## Supporting information for

**Supporting Figure 1.**
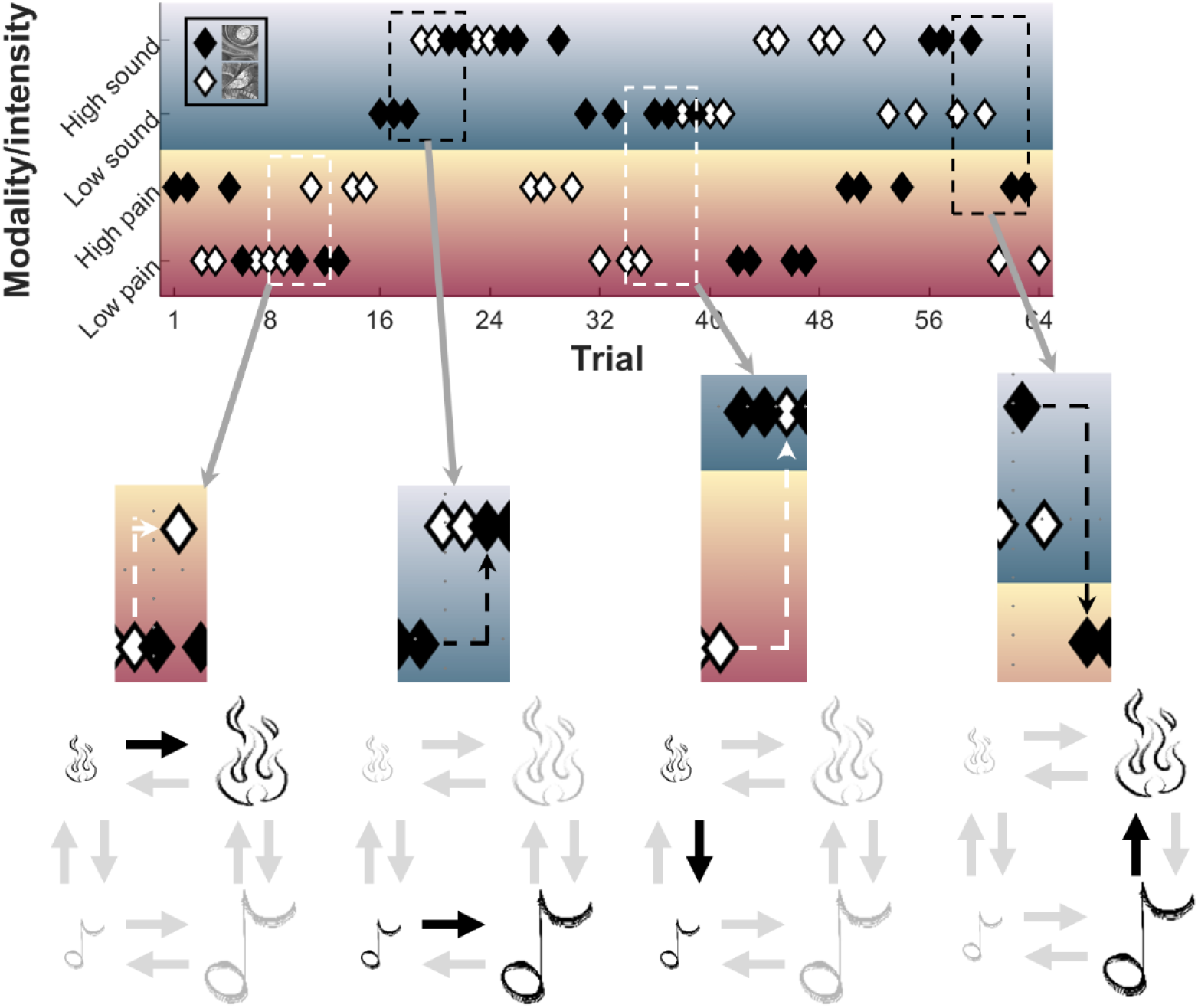
Illustration of reversal types. Both conditioned stimuli have an independent sequence of deterministic associations with one of the four unconditioned stimuli (also see Figure 2). The dashed lines illustrate reversals for CS1 (black) or CS2 (white). First column, CS2 intensity reversal from low to high heat; second column, CS1 intensity reversal from low to high sound; third column, CS2 modality reversal from low heat to low sound; fourth column, modality reversal from high sound to high heat.

**Supporting Figure 2.**
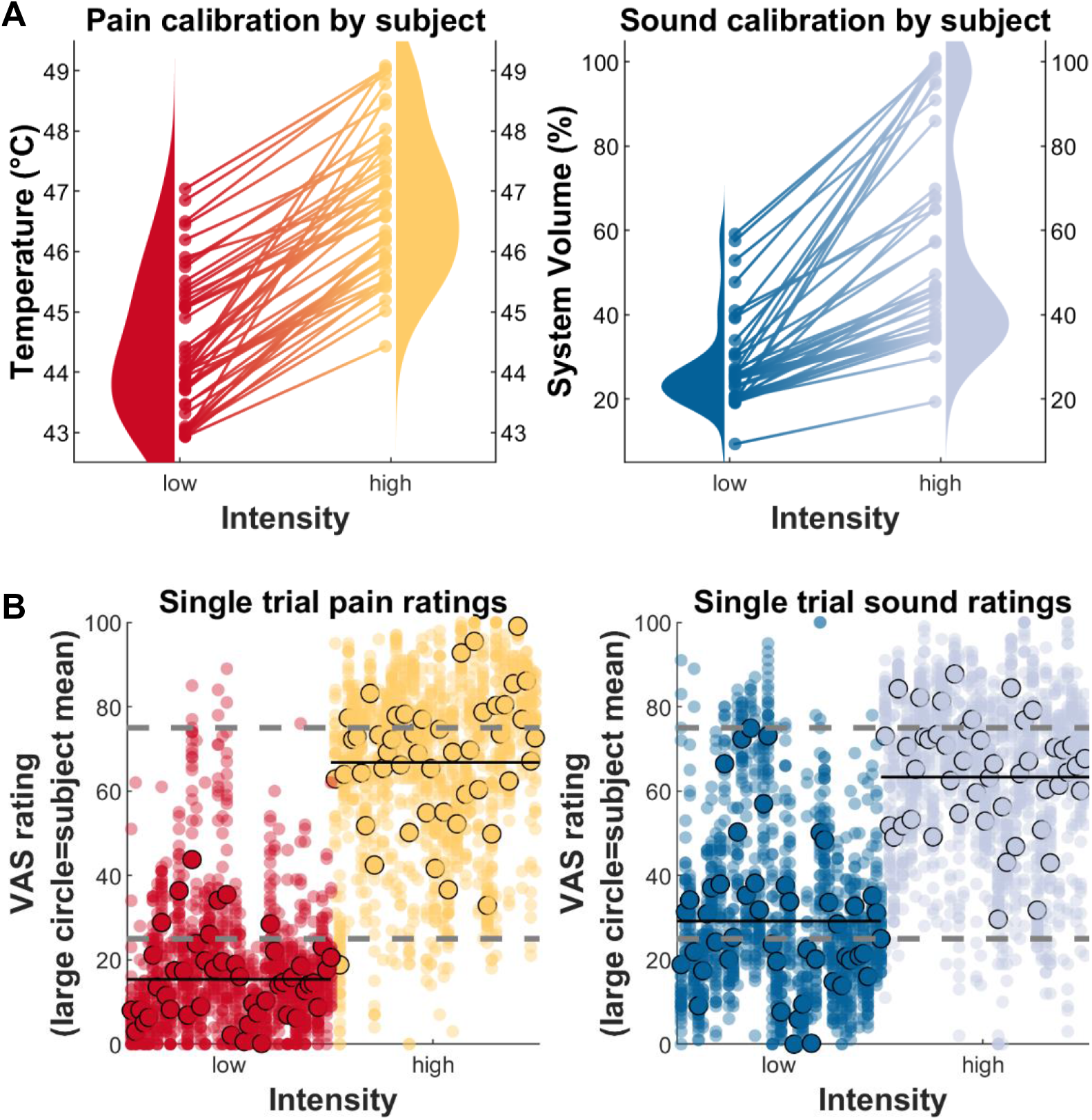
Behavioral results for low and high unconditioned pain and sound stimuli. **(A)** Calibrated stimulus intensities corresponding to VAS25 (low intensity) and VAS75 (high intensity) for pain stimuli and sound stimuli. Each line represents the two intensities per modality per subject; the violin plots aggregate over subjects. **(B)** Single trial ratings following pain stimulation and sound stimulation. Every column represents a single subject’s response to the respective intensity and modality; the bordered circle is a subject’s mean rating. The grey dashed lines is the “intended” rating as per calibration (VAS25 for low, VAS75 for high intensities). The black line is the actual mean rating over all subjects.

**Supporting Figure 3.**
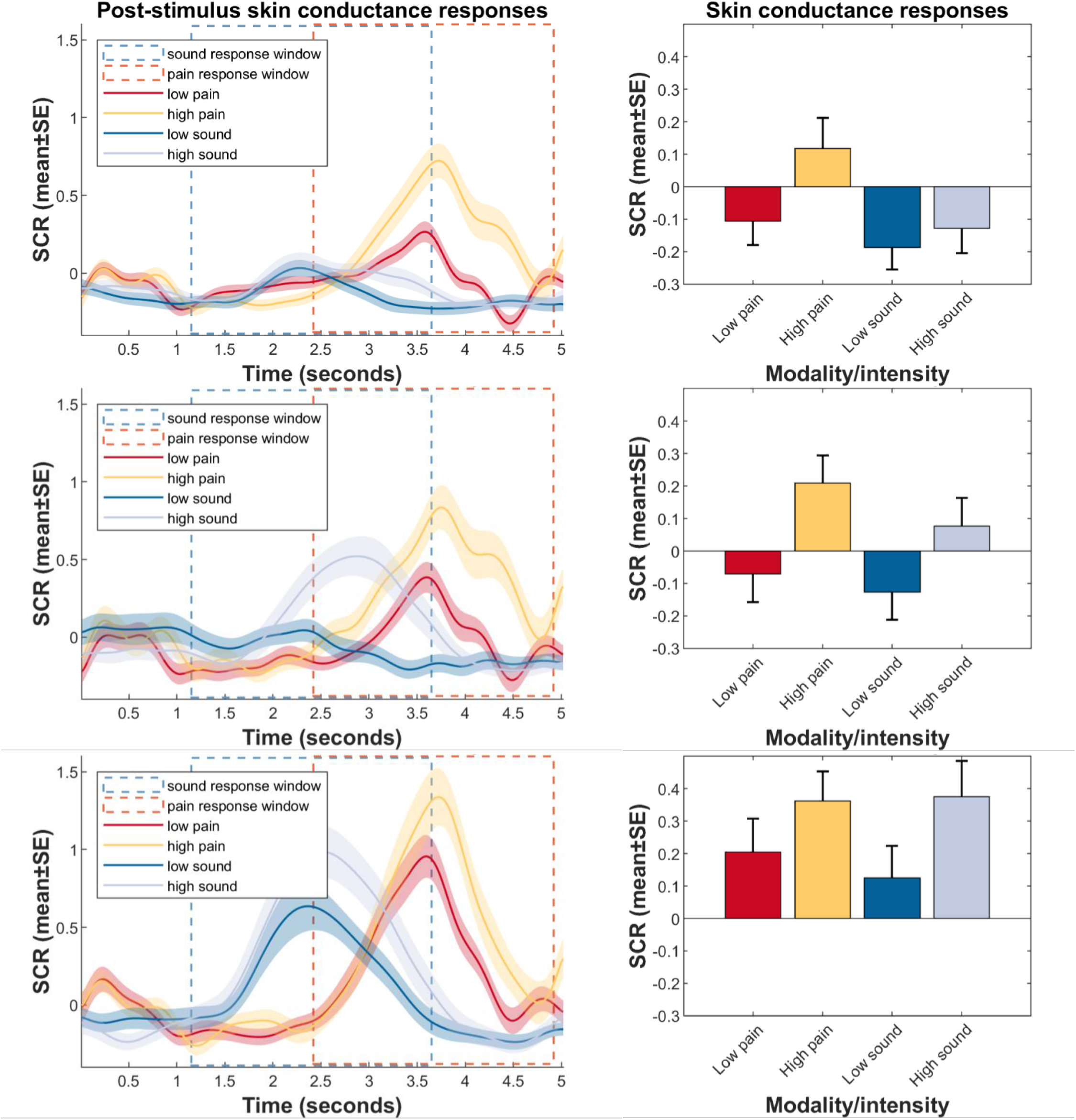
Results from skin conductance response measurements, by prediction error type. Rows show group means of SCR following no prediction error (row 1), intensity PE (row 2), and modality PE (row 3). Column show post-stimulus SCR (left) and SCR averaged within the indicated response windows (right).Differences between conditions are largest in the no PE condition, smallest in the modality PE condition, which also shows the largest SCR amplitudes. Statistics of differences between conditions are displayed in Supporting Table 1. All plots are based on log- and z-transformed data.

**Supporting Table 1.** Effects of modality and intensity, by prediction error type. Parameters obtained from linear mixed models with random subject intercept. Differences between the conditions are largest in trials with no prediction error, and smallest in trials with modality prediction error (cf. Supporting Figure 1).

| Subanalysis | Term | Estimate | SE | CI Lower | CI Upper | p |
| --- | --- | --- | --- | --- | --- | --- |
| No prediction error | Modality | -0.0442 | 0.0307 | -0.1044 | 0.0161 | 0.1506 |
|  | Intensity | 0.2444 | 0.0305 | 0.1845 | 0.3042 | 2 x 10<sup>-15</sup>* |
|  | Modality*Intensity | -0.2023 | 0.0433 | -0.2873 | -0.1173 | 3 x 10<sup>-6</sup>* |
| Intensity prediction error | Modality | -0.0506 | 0.0603 | -0.1689 | 0.0677 | 0.4014 |
|  | Intensity | 0.2479 | 0.0598 | 0.1305 | 0.3653 | 4 x 10<sup>-5</sup>* |
|  | Modality*Intensity | -0.026 | 0.0838 | -0.1904 | 0.1385 | 0.7566 |
| Modality prediction error | Modality | -0.056 | 0.0698 | -0.1929 | 0.0809 | 0.4227 |
|  | Intensity | 0.1138 | 0.071 | -0.0257 | 0.2532 | 0.1096 |
|  | Modality*Intensity | 0.0973 | 0.0994 | -0.0977 | 0.2924 | 0.3277 |

\*p < 0.001.

**Supporting Figure 4.**
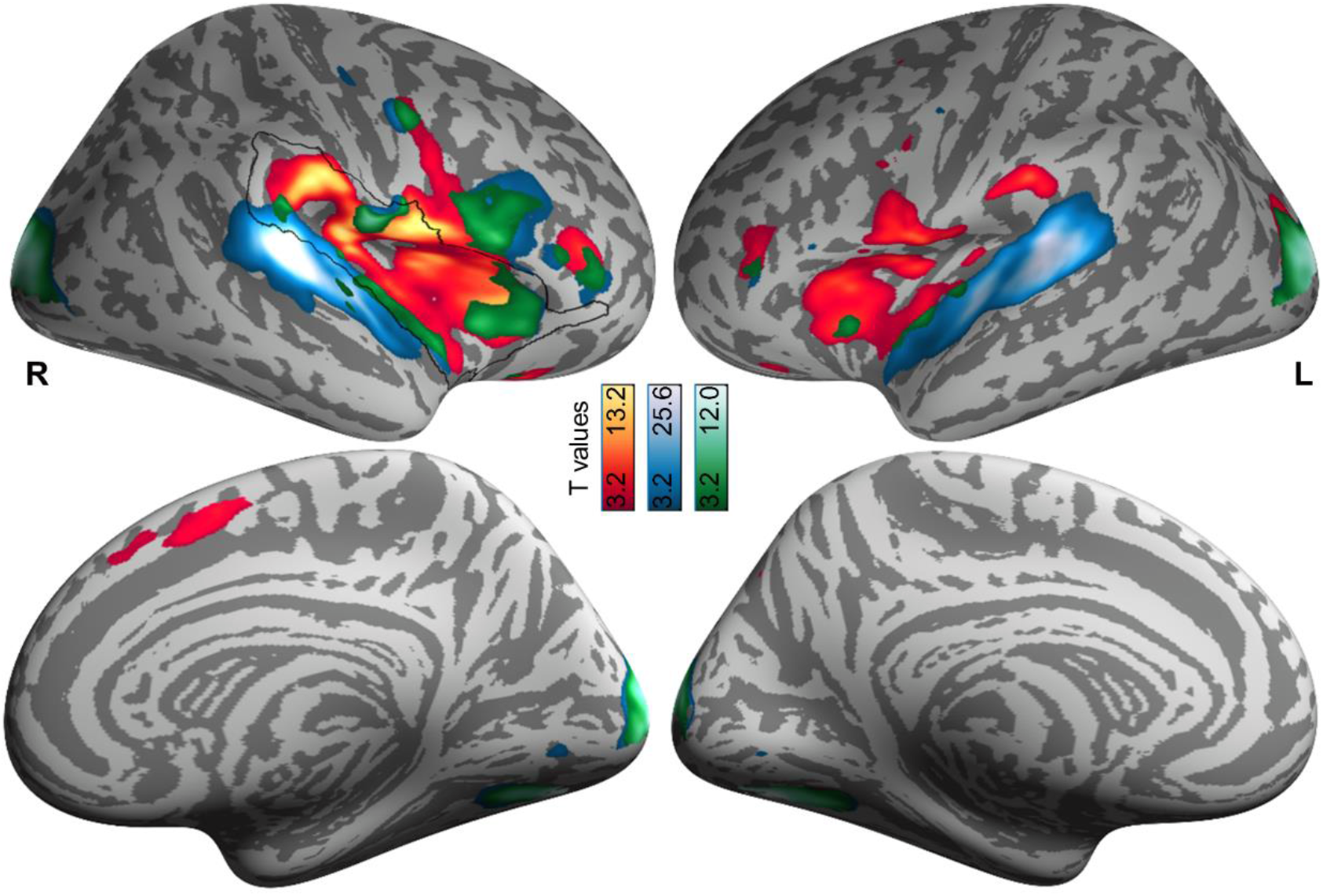
Lateral and medial views of brain surface results for heat onsets (yellow/red), sound onsets (blue), and their conjunction (green). Activations are overlaid on an average brain surface and thresholded at p[uncorr.] < 0.001. The black line delineates the region of interest whose results are highlighted in Figure 5a/b. R, right hemisphere; L, left hemisphere.

**Supporting Figure 5.**
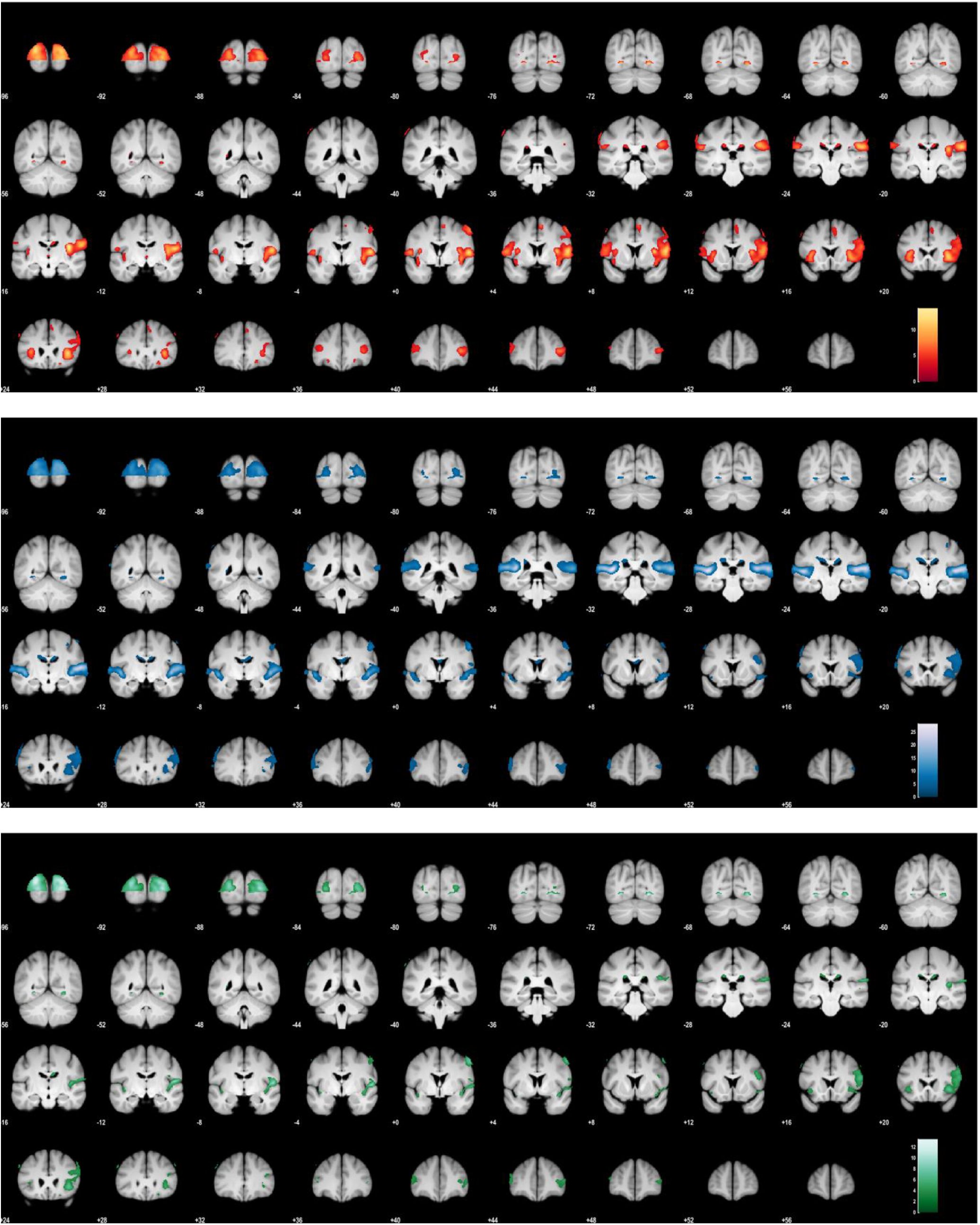
Brain volume results for heat onsets (yellow/red), sound onsets (blue), and their conjunction (green). Activations are overlaid on an average brain volume and thresholded at p[uncorr.] < 0.001.

**Supporting Figure 6.**
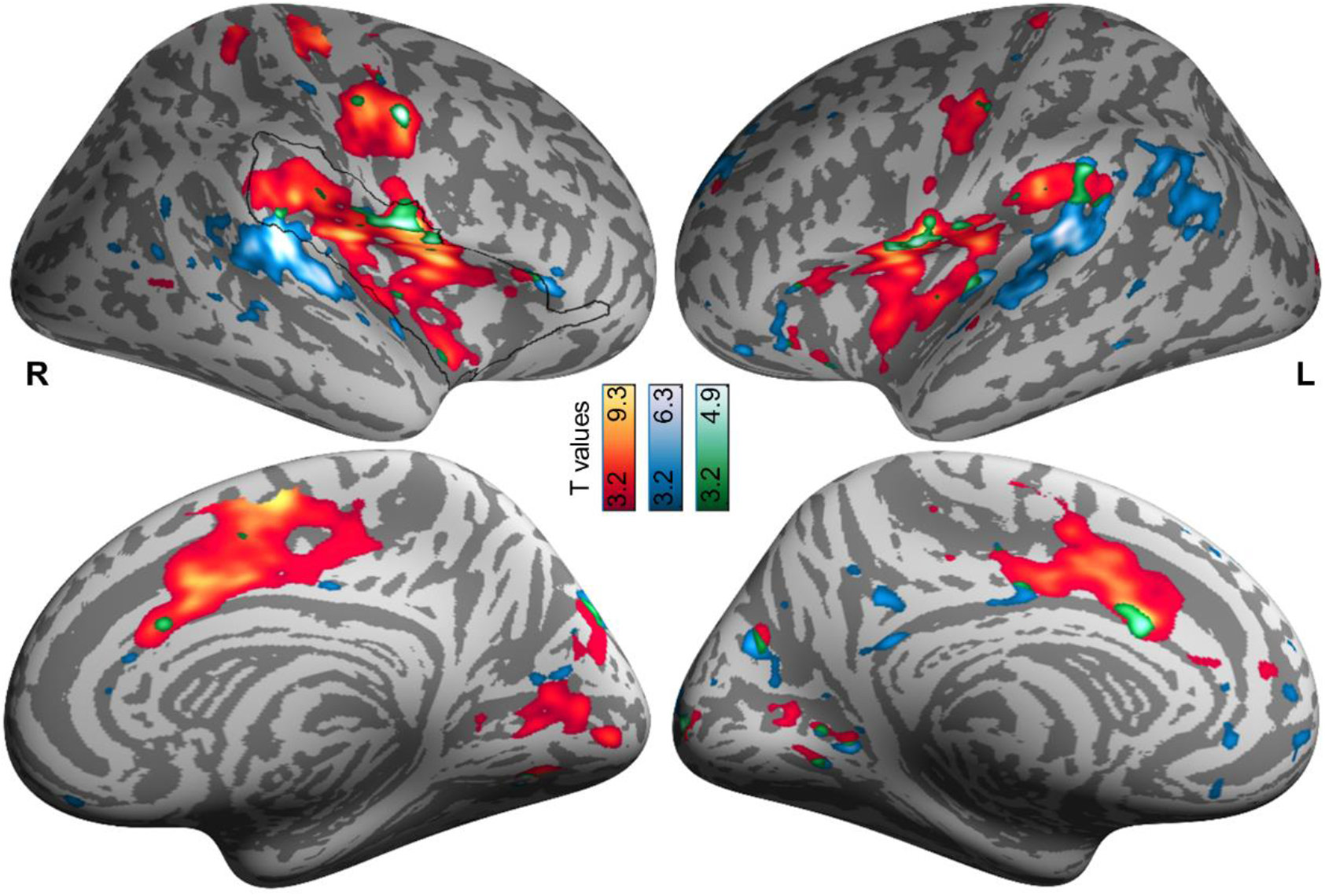
Lateral and medial views of brain surface results for pain ratings (yellow/red), sound ratings (blue), and their conjunction (green). Activations are overlaid on an average brain surface and thresholded at p[uncorr.] < 0.001. The black line delineates the region of interest whose results are highlighted in Figure 5b/c. R, right hemisphere; L, left hemisphere.

**Supporting Figure 7.**
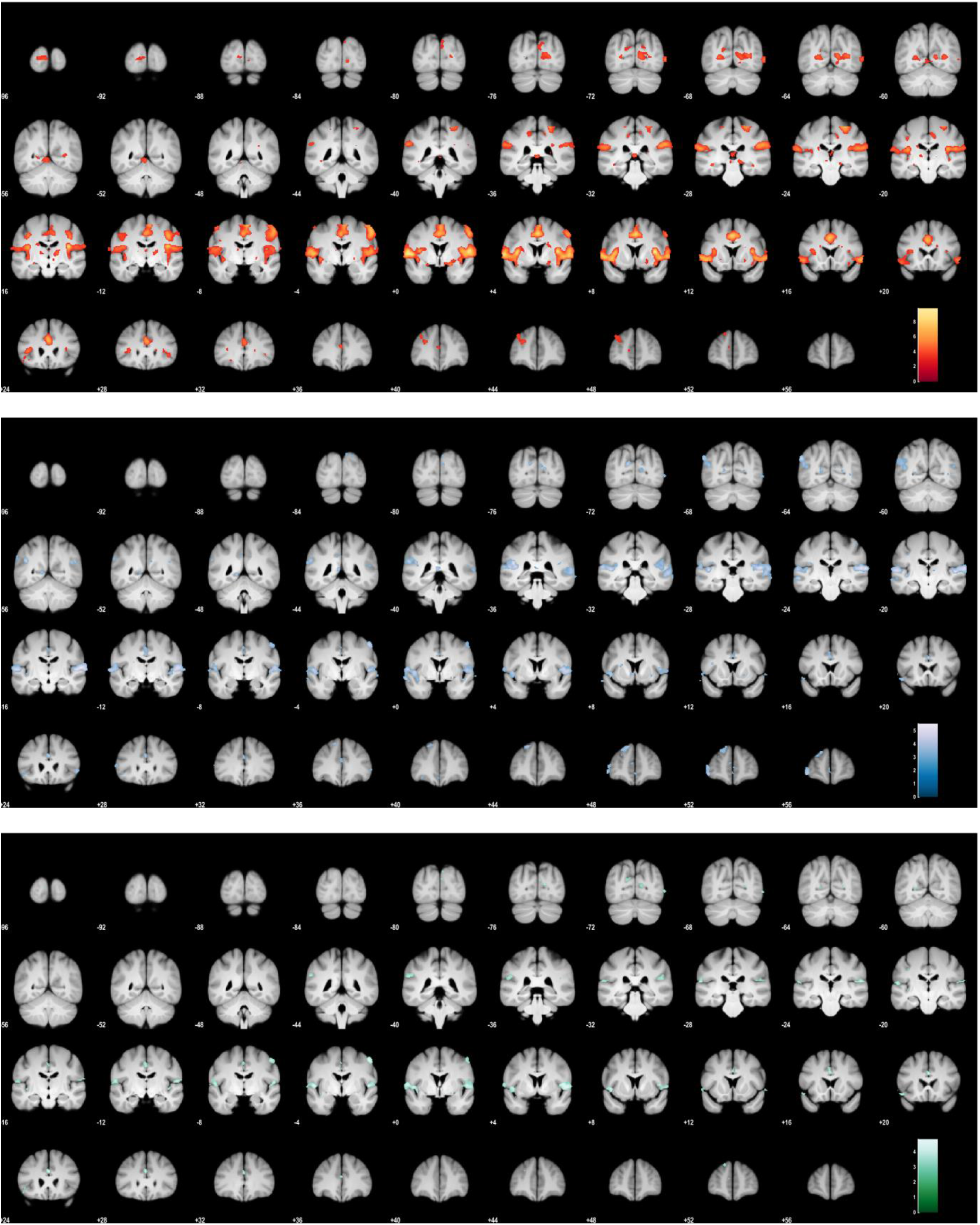
Brain volume results for pain ratings (yellow/red), sound ratings (blue), and their conjunction (green). Activations are overlaid on an average brain volume and thresholded at p[uncorr.] < 0.001.

**Supporting Figure 8.**
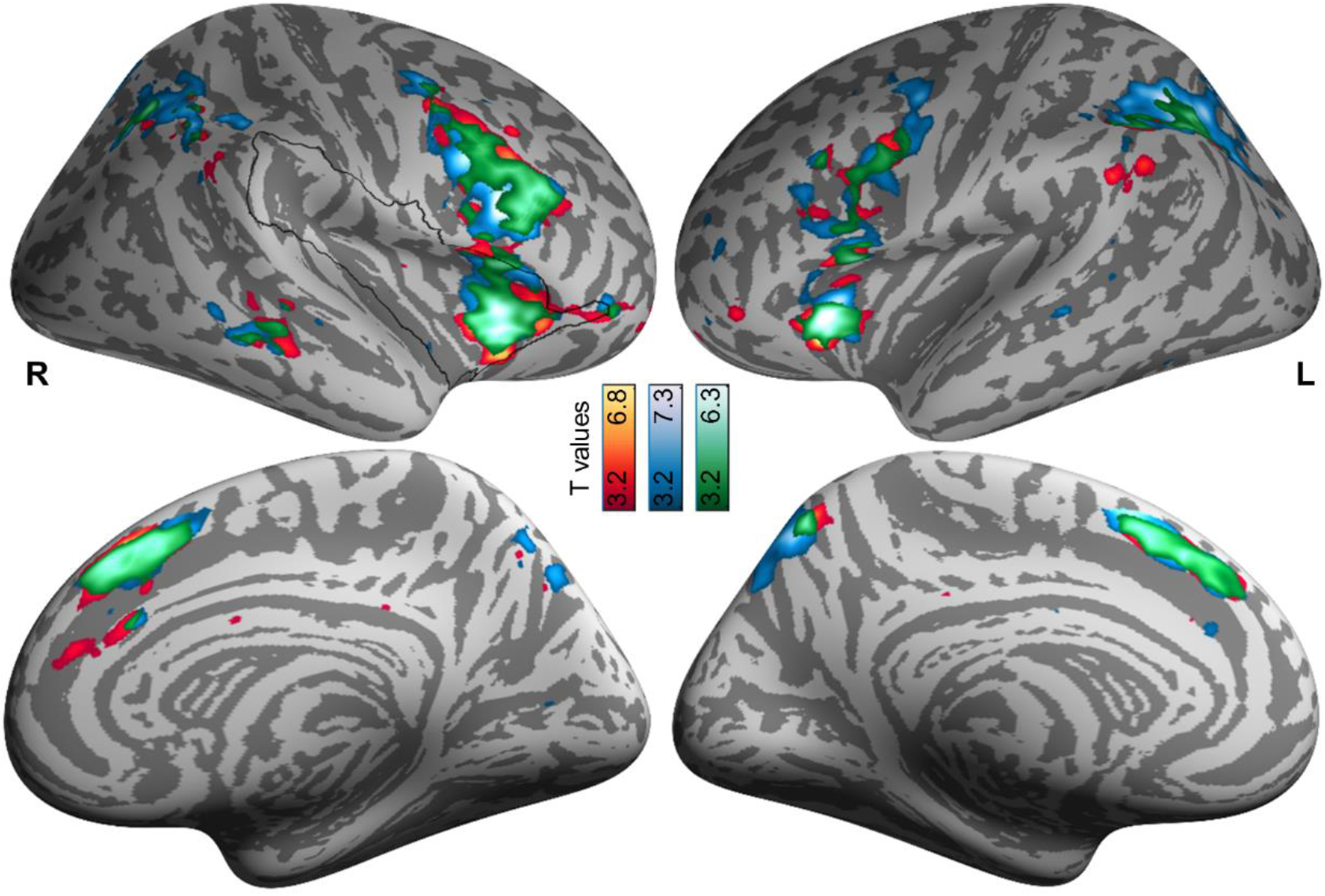
Lateral and medial views of brain surface results for unsigned intensity prediction errors for heat (yellow/red), sound (blue), and their conjunction (green). Activations are overlaid on an average brain surface and thresholded at p[uncorr.] < 0.001. The black line delineates the region of interest whose results are highlighted in Figure 6. R, right hemisphere; L, left hemisphere.

**Supporting Figure 9.**
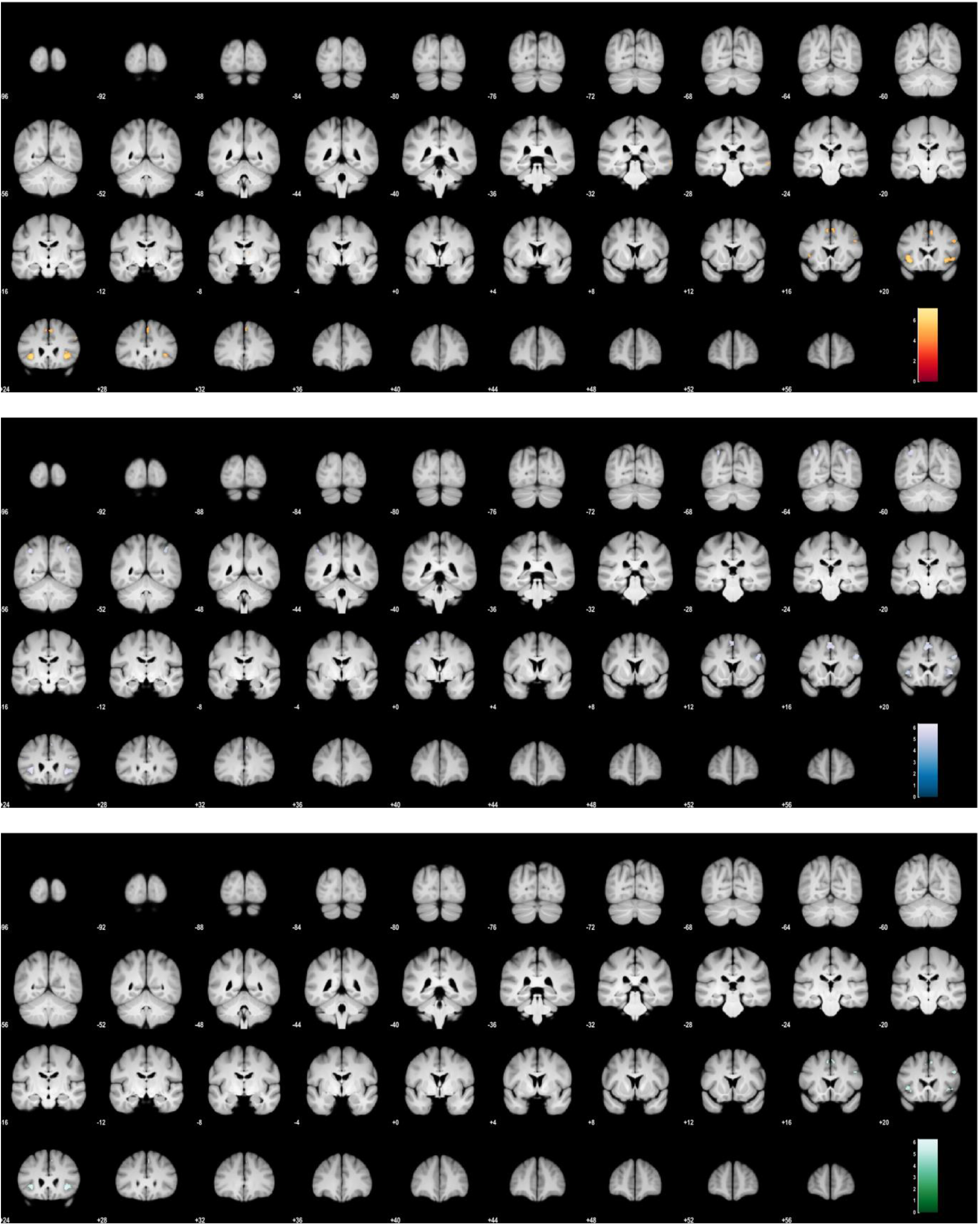
Brain volume results for unsigned intensity prediction errors for heat (yellow/red), sound (blue), and their conjunction (green). Activations are overlaid on an average brain volume and thresholded at p[uncorr.] < 0.001.

**Supporting Figure 10.**
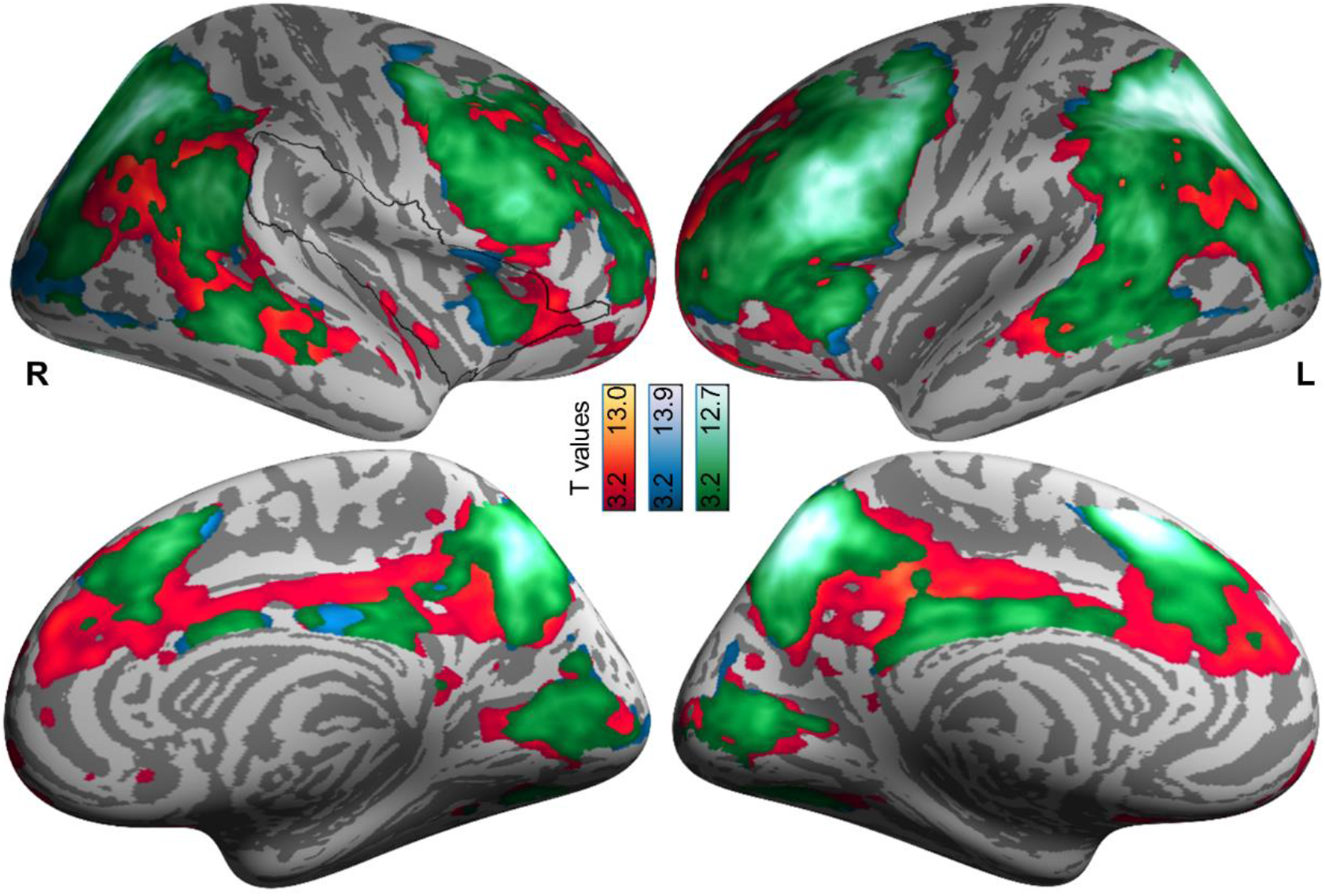
Lateral and medial views of brain surface results for modality prediction errors for heat (yellow/red), sound (blue), and their conjunction (green). Activations are overlaid on an average brain surface and thresholded at p[uncorr.] < 0.001. The black line delineates the region of interest whose results are highlighted in Figure 7. R, right hemisphere; L, left hemisphere.

**Supporting Figure 11.**
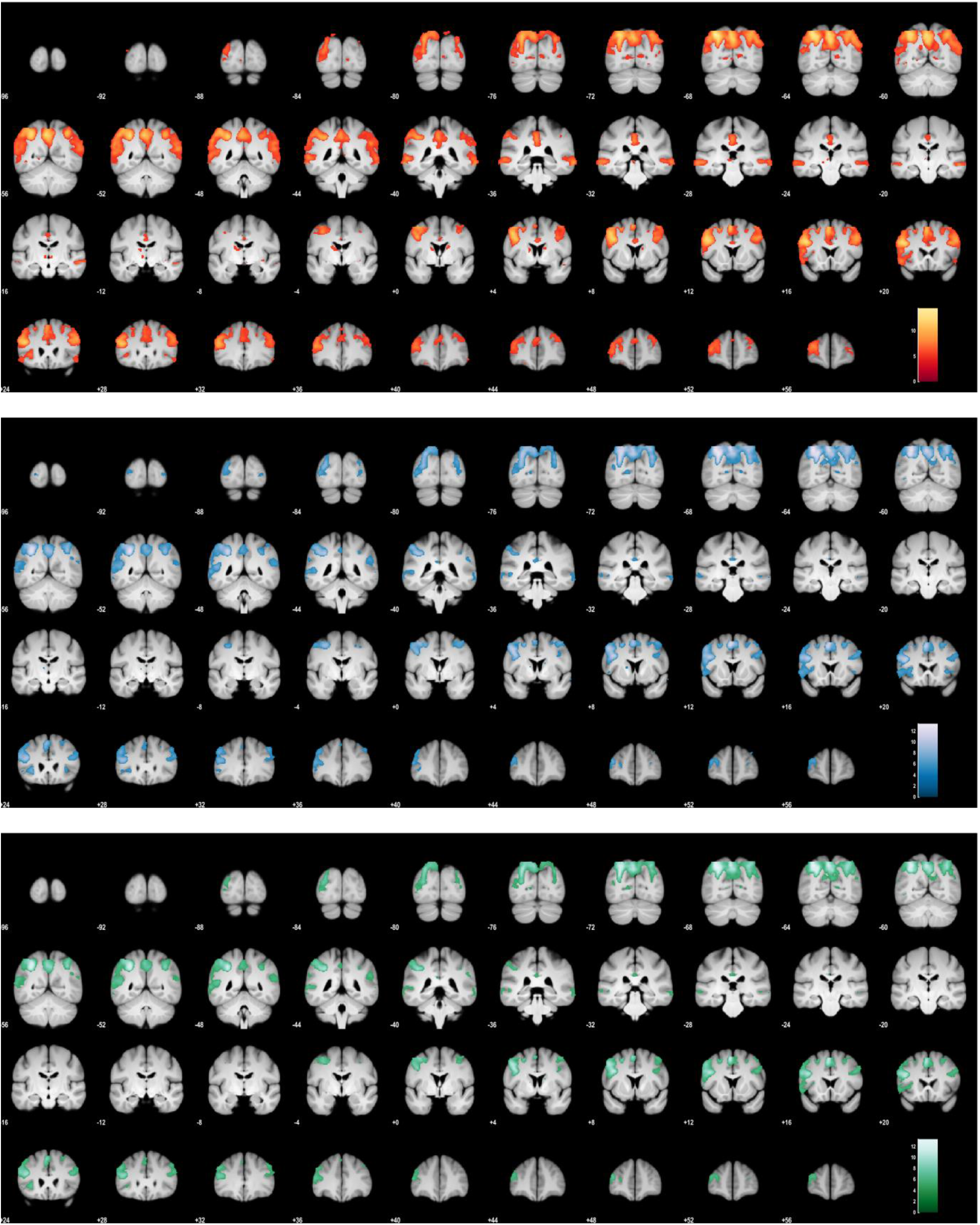
Brain volume results for modality prediction errors for heat (yellow/red), sound (blue), and their conjunction (green). Activations are overlaid on an average brain volume and thresholded at p[uncorr.] < 0.001.

**Supporting Figure 12.**
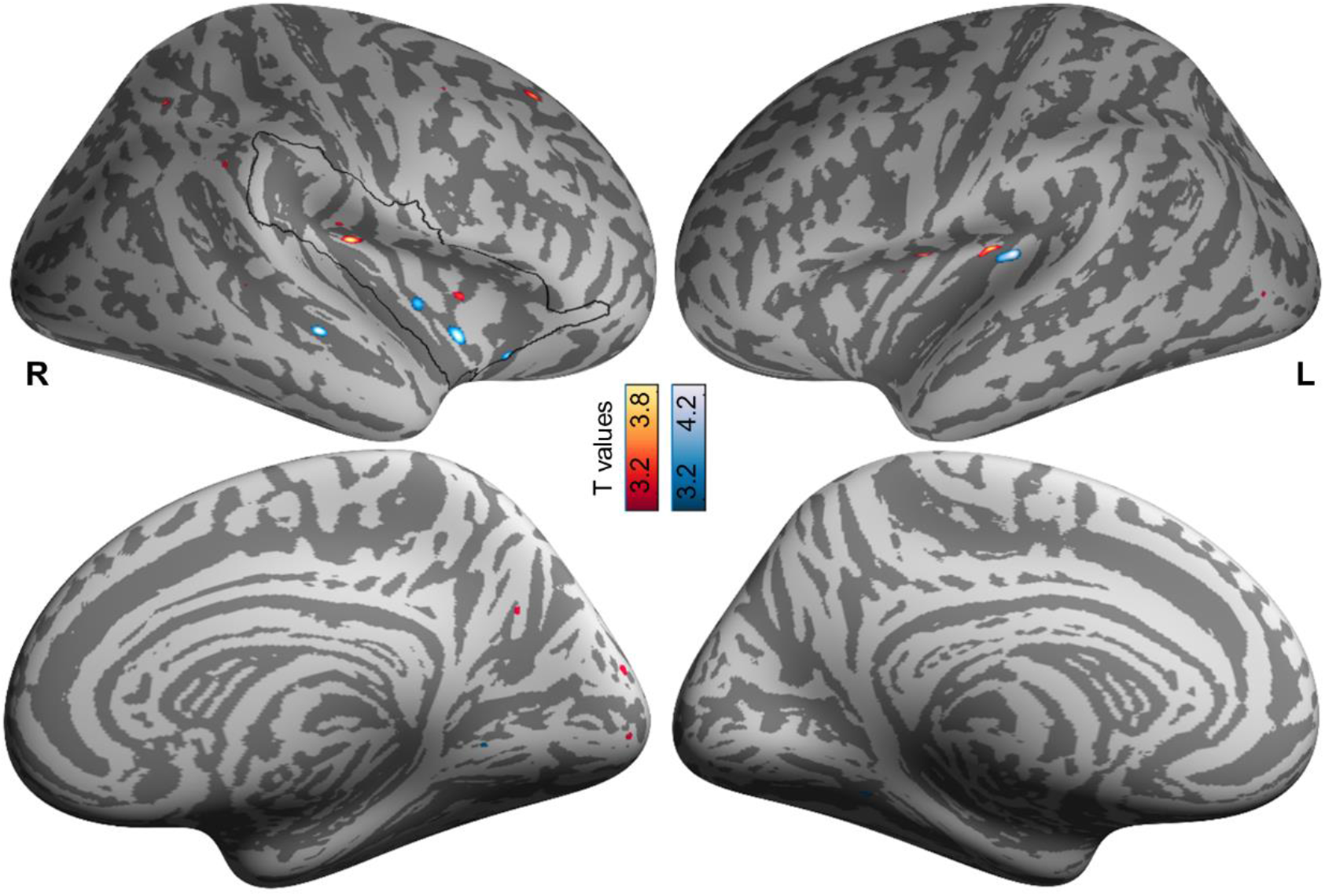
Lateral and medial views of brain surface results for signed intensity prediction errors for heat (yellow/red) and sound (blue). No significant conjunction activation prevails. Activations are overlaid on an average brain surface and thresholded at p[uncorr.] < 0.001. The black line delineates the region of interest whose results are highlighted in Figure 9. R, right hemisphere; L, left hemisphere.

**Supporting Figure 13.**
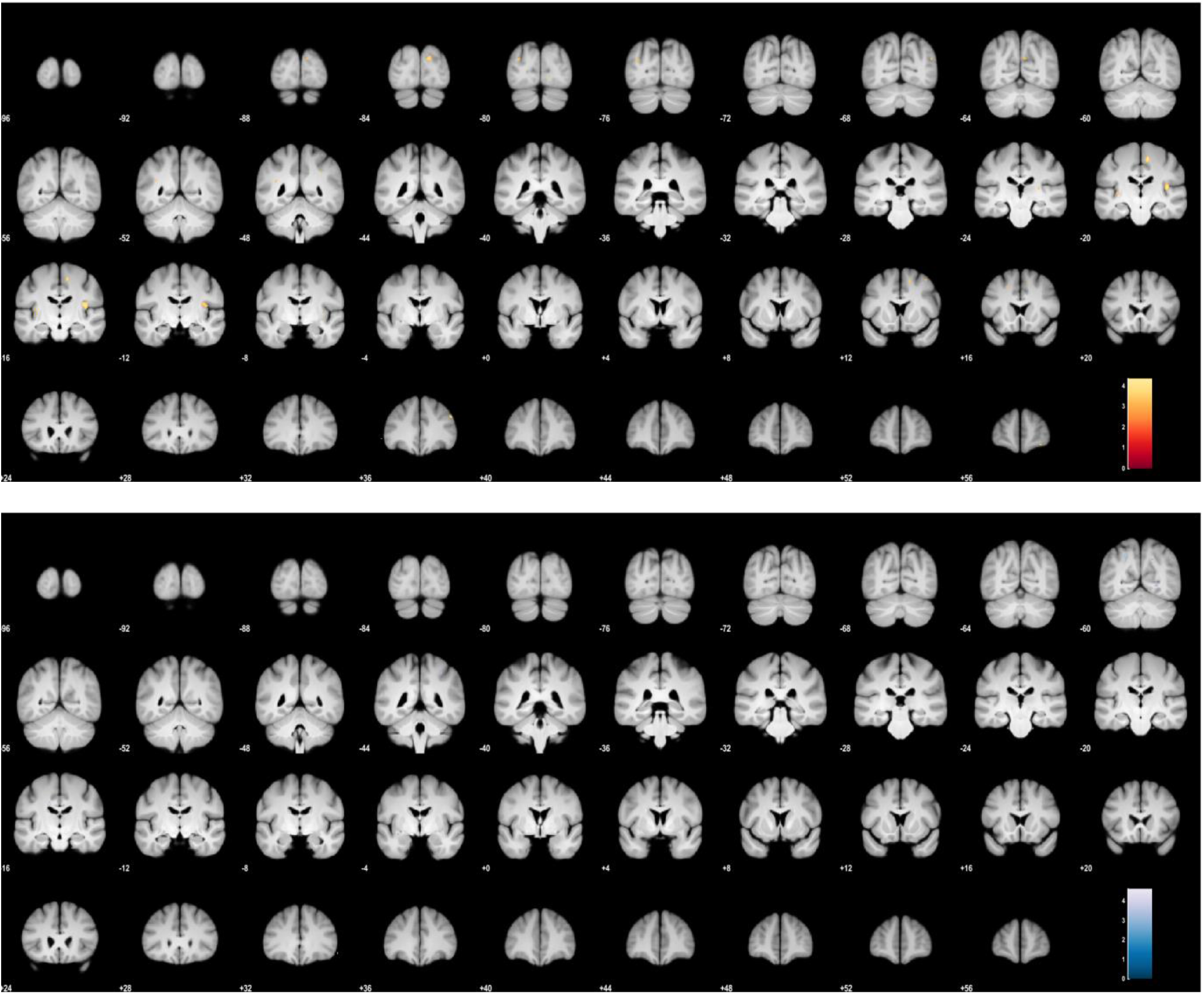
Brain volume results of signed intensity prediction errors for heat (yellow/red) and sound (blue). No significant conjunction activation prevails. Activations are overlaid on an average brain volume and thresholded at p[uncorr.] < 0.001.

**Supporting Figure 14.**
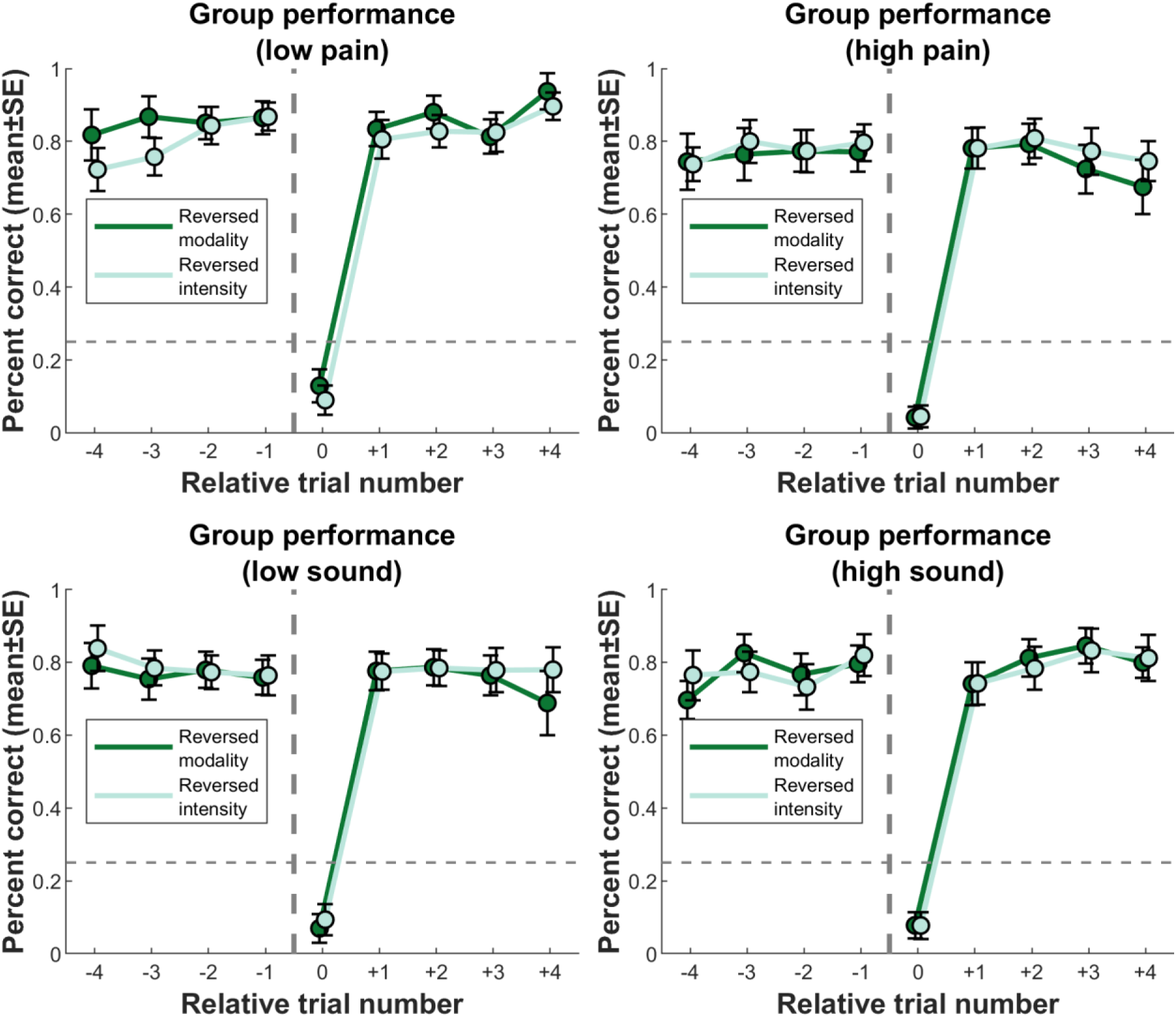
Mean performance split by modality/intensity. Grand mean performance is shown in Figure 3b.

**Supporting Table 2.** Sample characteristics. For references, see Supporting References.BDI-II, Beck Depression Inventory II; PHQ15, Patient Health Questionnaire-15; FPQ, Fear of Pain Questionnaire; PVAQ, Pain Vigilance and Awareness Questionnaire; PSQ, Pain Sensitivity Questionnaire; PRSS, Pain-Related Self-Statements; STAI, State-Trait Anxiety Inventory; MDMQ, Multidimensional Mood Questionnaire; exp., experiment.

| Questionnaire | Construct | Mean±SD | Sample range | Possible range |
| --- | --- | --- | --- | --- |
| **BDI-II** [SR 1,2] | Depression | 4.0±3.7 | 0-13 | 0-63 |
| **PHQ15** [SR 3] | Somatization | 3.4±2.7 | 0-10 | 0-30 |
| **FPQ** [SR 4] |  |  |  |  |
| severe | Fear of pain | 29.9±9.6 | 10-50 | 10-50 |
| minor | Fear of pain | 16.3±5.2 | 10-34 | 10-50 |
| **PVAQ** [SR 5] | Pain vigilance and awareness | 36.2±9.9 | 9-63 | 0-80 |
| **PSQ** [SR 6] | Pain sensitivity | 43.3±16.0 | 9-80 | 0-140 |
| **PRSS** [SR 7] |  |  |  |  |
| Catastrophizing | Pain catastrophizing | 8.4±6.1 | 1-27 | 0-45, higher more catastrophizing |
| Coping | Pain coping | 31.3±6.0 | 19-43 | 0-45, higher more active coping |
| **STAI** [SR 8,9] |  |  |  |  |
| Trait | Trait anxiety | 33.1±5.9 | 23-48 | 20-80 |
| State | State anxiety (pre experiment) | 33.4±6.2 | 23-49 | 20-80 |
| **MDMQ** [SR 10] |  |  |  |  |
| GoodBad A | Mood: Good vs bad (pre exp.) | 17.3±2.0 | 12-20 | 4-24, the higher the better mood |
| AwakeTired A | Mood: Awake vs tired (pre exp.) | 14.5±2.9 | 8-20 | 4-24, the higher the more awake |
| CalmNervous A | Mood: Calm vs nervous (pre exp.) | 16.1±2.2 | 10-20 | 4-24, the higher the calmer |
| GoodBad B | Mood: Good vs bad (post exp.) | 17.2±2.0 | 11-20 | 4-24, the higher the better mood |
| AwakeTired B | Mood: Awake vs tired (post exp.) | 11.7±3.1 | 7-18 | 4-24, the higher the more awake |
| CalmNervous B | Mood: Calm vs nervous (post exp.) | 17.2±2.4 | 11-20 | 4-24, the higher the calmer |

